# Cohesin bridging as a physical principle of enhancer-promoter communication

**DOI:** 10.64898/2026.05.07.723561

**Authors:** Timothy Földes, Karissa L. Hansen, Maxim Imakaev, Henrik D. Pinholt, Nezar Abdennur, Geoffrey Fudenberg, Irié Carel, Tayma Handal, Fernanda Vargas-Romero, Elphège P. Nora, Leonid A. Mirny

**Affiliations:** Massachusetts Institute of Technology, Institute for Medical Engineering and Science, Cambridge, MA, USA; Institut Curie, UMR3664 Nuclear Dynamics, Paris, France; Cardiovascular Research Institute, University of California, San Francisco, San Francisco, CA, USA; Developmental and Stem Cell Biology Graduate Program, University of California, San Francisco, San Francisco, CA, USA; Department of Genomics and Computational Biology, UMass Chan Medical School, Worcester, MA 01605, USA; Department of Systems Biology, UMass Chan Medical School, Worcester, MA 01605, USA; Department of Quantitative and Computational Biology, University of Southern California, Los Angeles, CA 90007, USA; Chan-Zuckerberg Biohub, San Francisco, CA, USA; Department of Biochemistry and Biophysics, University of California, San Francisco, San Francisco, CA, USA

## Abstract

Central to genome function, enhancers are non-coding sequences that can control transcription from promoters hundreds of kilobases away. Yet the physical basis of this long-range communication remains unclear. A prevalent view is that enhancers activate promoters when the two elements come into spatial proximity through the 3D folding of chromatin. However, activation by spatial proximity alone has struggled to explain several core features of enhancer function. Here, we propose that the molecular motor cohesin transmits long-range enhancer action by forming bridges between enhancers and promoters during loop extrusion. In this view, rare and transient bridges carry regulatory communication, rather than mere spatial proximity. We develop a quantitative model that predicts transcriptional output from cohesin-bridging dynamics and validate it by engineering cells in which strategically positioned CTCF sites rewire loop extrusion trajectories. The model explains how enhancer action scales with genomic distance, and how it can be either facilitated or insulated by CTCF sites across two orders of magnitude–behaviors incompatible with proximity-based models. Finally, our framework reveals that CTCF sites can block enhancers bidirectionally, by either blocking or releasing cohesin loops, resolving longstanding paradoxes between their effects on transcriptional regulation and genome folding. Together, our results establish cohesin bridging as a mode of enhancer-promoter communication that can be modulated by genomic context to achieve selective and tunable transcriptional control over long genomic distances.

## Introduction

Precise long-range transcriptional regulation is essential for developmental processes and its misregulation underlies a broad spectrum of human diseases. Yet, how genomic architecture translates into quantitative transcriptional outcomes remains unresolved–limiting both our mechanistic understanding of transcriptional regulation and our ability to interpret genetic variation. In vertebrates, long-range enhancer-promoter communication is understood to largely occur within topologically associating domains (TADs) (*1, 2*). DNA loop extrusion by the cohesin complex mediates the chromosomal interactions inside TADs, while CTCF binding sites delineate TAD boundaries (*3–8*) that restrict regulatory interactions (*9, 10*). Yet the physical basis of this long-range communication, and how enhancers selectively activate specific promoters within a genomic neighborhood, remain unclear (*11*).

A prevalent view is that enhancers activate promoters when the two elements come into spatial proximity. However, transcriptional activation by mere spatial proximity has struggled to explain several core features of enhancer function. First, enhancer activity does not directly track with the 3D contact frequency measured by chromosome conformation assays such as Hi-C, as exemplified by the fact that enhancer activity decays substantially more slowly with genomic distance than contact frequency. Second, TAD boundaries can strongly insulate promoters from neighboring enhancers despite causing only modest, ∼2-3-fold reductions in cross-boundary contacts (*1, 12–16*). Third, transcriptional dysregulation is challenging to predict from perturbations that alter 3D contacts: genomic manipulation of CTCF sites often produces effects that depend on the broader genomic context (*17–31*), while acute depletion of cohesin or CTCF rapidly disrupts TAD folding but cannot be sustained long enough to measure full transcriptional changes without incurring confounding secondary effects (*6, 7, 32–34*). Together, these observations make clear that 3D contact between enhancers and promoters does not simply equate to functional interaction. What mechanism then, if not mere spatial proximity, enables long-range enhancer–promoter communication?

Here, we consider the possibility that encounters bridged by cohesin represent a functionally privileged subset of enhancer-promoter contacts–distinct from diffusion-driven proximity–and explore how the unique dynamics of such bridges quantitatively regulate transcriptional outcomes. We formalized this hypothesis in a quantitative framework that links cohesin-bridging to transcriptional output, and generated explicit predictions for how the genomic arrangement of an enhancer, a promoter, and CTCF sites can tune transcriptional outcomes. Genome-engineering experiments quantitatively validate these predictions, establishing cohesin bridging as a physical principle for enhancer-promoter communication. This model accurately predicts the quantitative effects of genomic distance on enhancer action, unifies the contrasting effects of CTCF as transcriptional insulator and facilitator as manifestations of the same underlying physical principle, and resolves long-standing paradoxes between how three-dimensional proximity relates to enhancer action. Beyond illuminating fundamental principles of enhancer-promoter communication, this work provides a foundation for interpreting the effects of structural variants and for designing new synthetic regulatory architectures.

## Model

### Cohesin bridging displays distinct properties from 3D contacts

Cohesin loop extrusion can bring enhancers and promoters together either by compacting chromosome domains, which reduces the 3D search space for diffusion-driven encounters, or by creating direct bridges between the translocating loop bases (**Fig. 1A**). These two types of contacts differ fundamentally in their biophysical properties, creating distinct expectations for how they might relate to enhancer-promoter communication. First, diffusion in 3D is an undirected search process (**Fig. 1B**, top), with contact probabilities decaying rapidly as genomic distance increases (*16, 26, 35*), whereas loop extrusion reduces search to a one-dimensional scan that can remain processive over hundreds of kilobases (kb) (*3*). This makes cohesin especially important for distal, rather than proximal, enhancer-promoter communication (*21, 33, 36–40*). Second, spatial contacts only last on the order of seconds, while cohesin bridges can persist for tens of seconds up to minutes (*41–44*) (**Fig. 1B**, middle), offering a longer window for time-gated processes during initiation of transcription (*45*) (**Fig. 1B**, bottom). We therefore sought to derive a quantitative understanding of the transcriptional behaviors that specifically arise from cohesin-bridged encounters, as opposed to any possible type of 3D contact.

**Figure 1.**
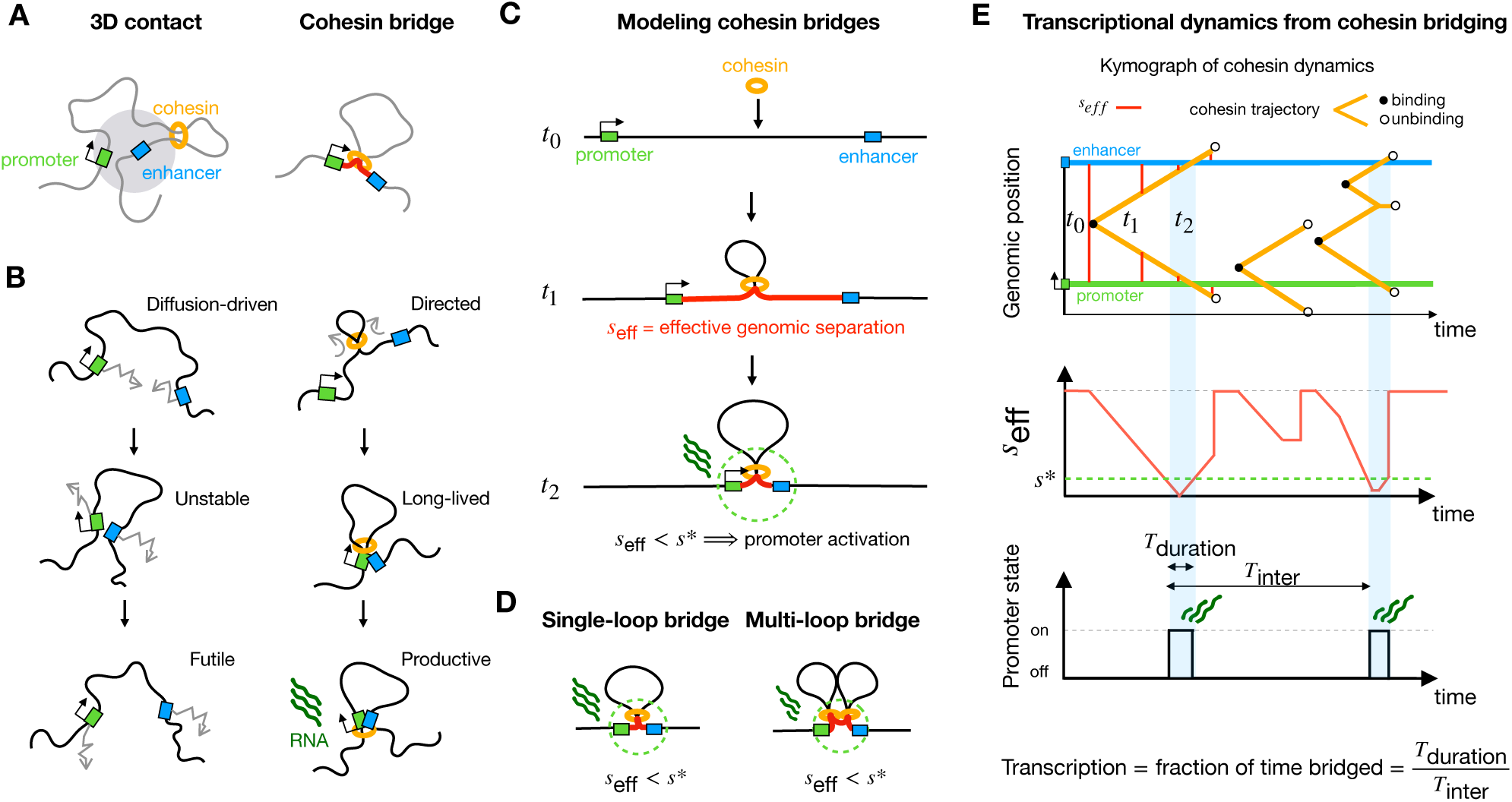
Modeling cohesin-bridged enhancer-promoter communication. **(A)** Comparison of diffusion-driven and cohesin-bridged enhancer-promoter contacts. **(B)** Enhancer-promoter contacts arising from random diffusion are non-directed, transient, and less likely to produce transcriptionally productive interactions (left). Cohesin loop extrusion generates directed, longer-lived contacts that offer longer support to transcription initiation (right). **(C)** Cohesin translocation dynamics are modeled on a one-dimensional lattice. As the cohesin loop grows, the effective separation (*s_eff_*) between the enhancer and promoter shrinks. Promoter activation is considered to occur when *s_eff_* is smaller than the distance threshold for activation *s\**, which corresponds to the cohesin-bridged state. **(D)** Cohesin-bridging can occur through single-or multi-loop configurations. **(E)** Schematics depicting the transcriptional dynamics arising from cohesin bridges. Top, cohesin trajectories along the chromatin fiber and promoter/enhancer positions over time. Three events are illustrated: a single-loop contact that either bridges (left) or fails to bridge (center), and a multi-loop contact in which two cohesins cooperatively bridge (right). The resulting changes of *s_eff_* across time (middle) and promoter activation states (bottom) demonstrate how transcriptional output is determined.

### Modeling cohesin-bridged enhancer-promoter communication

Rather than modeling the full 3D dynamics of a fluctuating chromatin fiber, we sought to reduce the system to a one-dimensional description of cohesin extrusion, thereby isolating the specific contribution of cohesin bridging from other sources of enhancer-promoter proximity. We therefore represent the chromatin fiber as a one-dimensional lattice containing an enhancer and a promoter separated by a genomic distance 𝐿. Along this DNA segment, cohesin loads uniformly (spaced on average by distance 𝑑) and extrudes at constant velocity *v* with a residence time 𝜏 (exponentially distributed), resulting in a processivity 𝜆 = *v*𝜏, i.e. the average loop size for an unobstructed cohesin (**fig. S1, A** and **B, Methods**). Cohesins block each other upon colliding while continuing to extrude on unblocked sides. A CTCF site is modeled as an extrusion barrier that blocks cohesins approaching its front side with probability 𝑝_!_while allowing cohesins approaching from the back side to bypass (see Methods), unless stated otherwise.

As cohesin extrudes a loop, it shortens the effective separation *s_eff_* between the enhancer and the promoter (**Fig. 1C**). An enhancer-promoter bridge is considered to occur for as long as *s_eff_* is below the genomic separation threshold for activation *s\**. The bridge may be mediated by a single cohesin loop or by multi-loops created from stacked cohesins (**Fig. 1D**). The threshold *s\** therefore represents the separation below which an enhancer activates the promoter without requiring further extrusion. Maintaining the enhancer and promoter at a separation of *s\** (or less) provides an opportunity for multiple molecular interactions during the lifetime of the bridge (*s_eff_* <*s\**). The value of *s\** may depend on promoter sensitivity and enhancer strength, with weaker elements possibly requiring closer proximity (smaller *s\**).

When such a bridge is formed, we consider the promoter to switch into an active state, from which transcription occurs at a constant rate (Methods). The total transcriptional output is then determined by the total fraction of time in the bridged state, i.e. the product of bridge frequency (1/*T_inter_*, with *T_inter_* being the time between bridge onset) and duration (*T_duration_*) (**Fig. 1E and fig. S1C**).

This approach enables modeling cohesin dynamics and the resulting transcriptional output for different enhancer, promoter, and CTCF arrangements, both analytically and through simulations (see Methods). The analytical theory treats cohesins as non-interacting, neglecting collisions between neighboring cohesins and the contribution of multi-loop bridges, but provides intuition for how biophysical parameters (cohesin residence time, velocity, density, etc.) affect bridging kinetics and transcriptional output. Simulations explicitly model cohesin collisions, capturing multi-loop configurations that arise when neighboring cohesins collide, at the cost of requiring numerical computation. We provide a video to illustrate the key features of the cohesin bridging model through multiple linked visualizations of simulated trajectories (Video S1). Together, this approach yielded explicit transcriptional predictions that we tested experimentally by creating a series of genome modifications that perturb enhancer-promoter cohesin bridges.

## Results

### The probability of enhancer-promoter bridging decays exponentially with genomic distance

In the cohesin-bridging model, the search process between the enhancer and the promoter requires two steps: first, cohesin loads on chromatin; second, cohesin must reach both regulatory elements without dissociating.

For a single cohesin to eventually bridge the enhancer and the promoter, it must load in a specific location we refer to as the enhancer-promoter *antenna* (**Fig. 2A**), by analogy to other search processes on DNA (*46–48*). For symmetric two-sided extrusion, cohesins loading outside this antenna would extrude the enhancer and promoter asynchronously, never directly bridging them (**Fig. 2A**), and therefore making no direct contribution to transcriptional activation.

**Figure 2.**
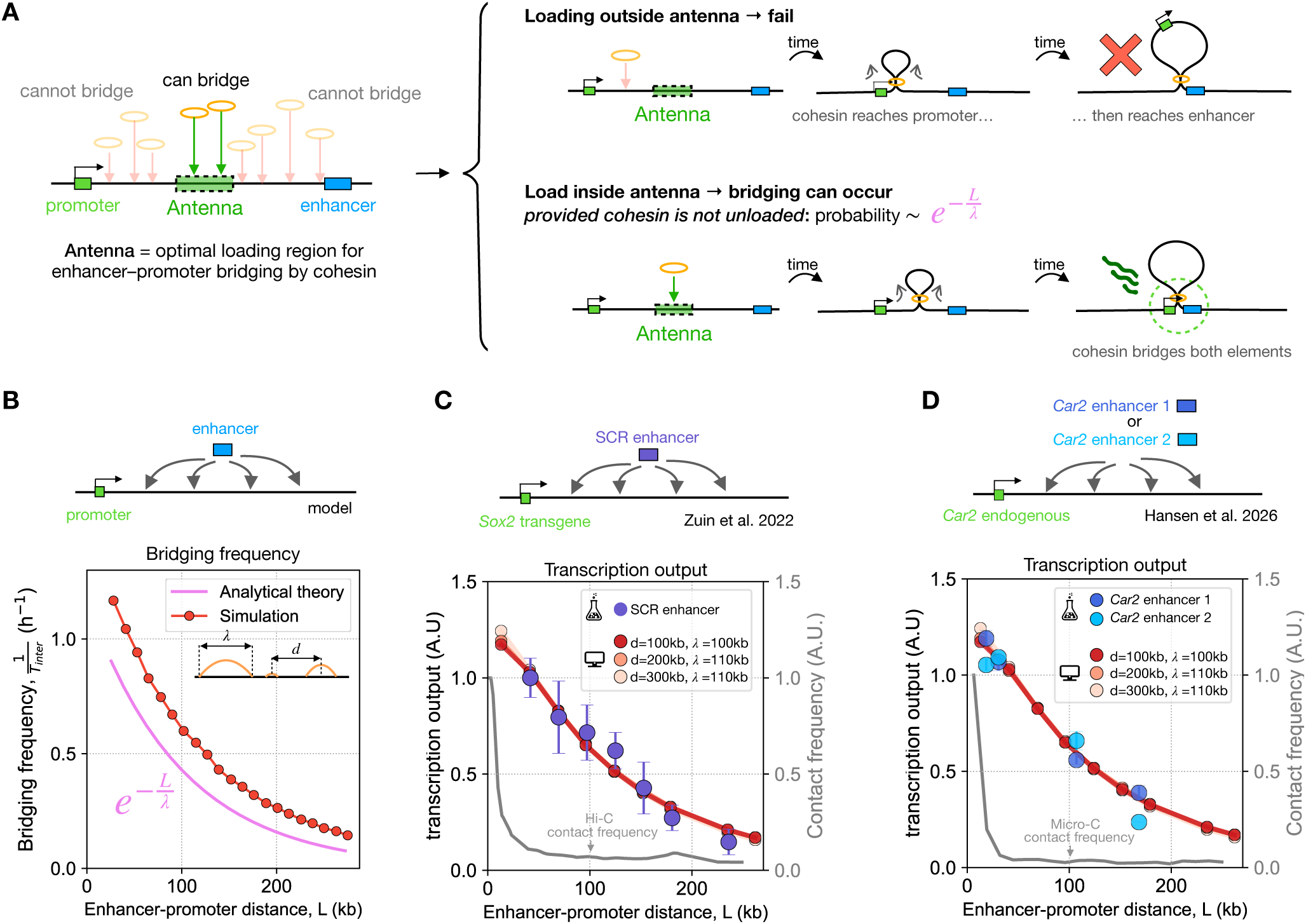
Regulatory communication through cohesin bridging explains how enhancer action scales with genomic distance. **(A)** The antenna defines the genomic region from which cohesin loading is most likely to create an enhancer-promoter bridge. For symmetric extrusion the antenna is around the mid-point of the enhancer-promoter interval (shown). Cohesin loading outside the antenna fails to bridge the enhancer with the promoter, as cohesin reaches each element asynchronously. A cohesin that is loaded within the antenna can reach both elements simultaneously, provided it does not unload before. The probability that a cohesin loaded within the antenna extrudes to create an enhancer-promoter bridge is ∼ 𝑒 ^“#/%^, where *L* is the enhancer-promoter genomic distance and 𝜆 is cohesin processivity. **(B)** Exponential decay of cohesin-bridging probability from simulations and analytical theory. Analytical formula is given in methods and is derived in the supplementary text. Inset: schematic of key model parameters: 𝜆 and the average distance between cohesins *d*. **(C)** The cohesin bridging model properly captures the transcriptional output measured from a transgenic *Sox2* promoter as a function of genomic distance to insertions of the SCR enhancer (*26*). Curves show best-fitting simulations for a range of cohesin separation (*d*) and cohesin processivity (𝜆; see fig. S4 for fitting details). Experimental values correspond to binned averages of individual cell lines grouped by genomic distance (27 kb bins; mean ± SEM). Transcriptional output (A.U.) was normalized to highlight scaling over absolute enhancer strength. The grey curve corresponds to the Hi-C contact frequency between the *Sox2* transgene and the TAD into which it was inserted (data from (*26*)), which decays much more steeply than transcription. **(D)** The same model parameters recapitulates the distance-dependent decay of transcription measured at the endogenous *Car2* locus (*33*), without further fitting. Transcriptional output (A.U.) was normalized as in (C). The grey curve corresponds to the Micro-C contact frequency between the *Car2* promoter and its TAD (data from (*74*)).

For symmetric two-sided extrusion, the antenna is centered at the midpoint of the enhancer-promoter interval, and its length 𝑙 is approximately equal *s\** (plus the lengths of the enhancer and promoter, which are typically much shorter; **fig. S2, A and B**). For other extrusion dynamics, such as extrusion with direction switching, the antenna becomes more diffuse, but the fraction of cohesins that can successfully bridge remains constant (**fig. S2, C and D**).

Assuming a constant dissociation rate yields an exponentially decreasing probability of bridging as genomic distance increases. Specifically, the probability that cohesins with an average processivity 𝜆 survive to bridge an enhancer and promoter separated by a genomic distance 𝐿 is given by 𝑒^“#/%^. When enhancer-promoter separation greatly exceeds cohesin processivity, direct bridging by a single cohesin becomes unlikely. However, multiple collided cohesins can jointly bridge the intervening distance.

Such multi-loop configurations constitute the predominant mode of enhancer-promoter bridging at large separations, and enable cohesins loaded outside the antenna to indirectly contribute to bridging (**fig. S3**).

### Enhancer action scales exponentially with genomic distance

Analytical theory (see supplementary text for derivation) and simulations thus show that the frequency of cohesin bridges between the enhancer and the promoter decays exponentially as their genomic separation increases, which is in sharp contrast to the power-law decay of 3D contact frequency (*35, 49*) (**Fig. 2C-D**).

We therefore asked to what extent distance-dependent transcriptional activity in experiments (*26, 33*) can be quantitatively described by cohesin bridging. First, we fitted a dataset previously obtained by transposing the *Sox2* control region (SCR) enhancer up to ∼250 kb around a transgenic *Sox2* promoter in mouse embryonic stem (ES) cells (*26*). Sweeping over a broad range of model parameters (cohesin processivity 𝜆, distance between cohesins 𝑑, and genomic separation threshold for bridging *s\**), we identified cohesin-bridging parameters that reproduced the measured transcriptional outputs with remarkable accuracy (**Fig. 2C**). Best fits were largely insensitive to the specific values of 𝑑 or *s\** and clustered around 𝜆 ∼110 kb (**fig. S4, A and B**), in the range of independent estimates from Hi-C (*50, 51*). Moreover, using these parameters successfully reproduced independent experimental measurements of another prior study that relocated the *Car2* enhancers 1 and 2 (CE1 and CE2) around the endogenous *Car2* promoter (**Fig. 2D**) in mouse ES cells (*33*). Cohesin bridging thus provides a mechanism for long-range regulatory communication that decouples transcriptional output from 3D contact frequency, explaining how transcription can decay more gradually with enhancer-promoter genomic distance than 3D contacts (**Fig. 2C-D**).

Beyond predicting steady-state transcription, the model also describes the temporal dynamics of cohesin bridges. Using the same parameter range, we computed the inter-bridge times (*T_inter_*) and compared them to the inter-burst intervals experimentally measured for L=150 kb with a *Sox2* transcriptional reporter in ES cells (*52*) (**fig. S4C**). For cohesin residence time of 400 seconds and estimated processivity of 𝜆=100 kb, the model reproduced the observed inter-burst interval of ∼3 hours (*52*). This narrowed model parameters to 𝑑 ∼100-200 kb for the separation between cohesins, in the range recently estimated (*53–55*), and to *s\** ∼10-20 kb (genomic distance below which enhancer action becomes independent of cohesin, consistent with recent measurements (*33*)).

The scaling of enhancer action with genomic distance emerges as a direct consequence of extrusion dynamics, with no additional assumptions. This parsimony contrasts with previous models that invoked nonlinear promoter responses to explain how small changes in contact frequency can translate into large transcriptional effects (*26, 56*).

Since comprehensive catalogs of genetically validated enhancer-promoter pairs are not available we cannot immediately explore how this mechanism accounts for enhancer action genome-wide, where additional mechanisms beyond cohesin are also at play (*33, 57–60*). We therefore turned to a well-characterized locus in which distal enhancer action can be experimentally reduced to a single enhancer-promoter pair, and tested predictions of the bridging model by predictably altering cohesin trajectories through manipulation of CTCF binding sites.

### Biophysical basis for regulatory facilitation by flanking CTCF barriers

#### Transcriptional facilitation by CTCF at the *Car2* locus

CTCF binding sites are barriers to cohesin extrusion (*3, 4, 7*), accounting for both the enhancer-blocking insulator effect of CTCF (*61*) and the correspondence between TADs and *cis*-regulatory domains (*9, 10*). Yet CTCF can also facilitate enhancer-promoter communication when binding occurs directly flanking the enhancer or promoter (*24, 37, 62–64*). Whether insulation and facilitation reflect distinct properties of CTCF or are two manifestations of the same cohesin-based mechanism remains unclear.

To address this, we experimentally inserted CTCF sites at the *Car2* locus in ES cells, where distal enhancers are essential for transcriptional activation and rely on cohesin extrusion **(fig. S5, A to D)** (*33*). To simplify interpretations, we used a previously published cell line in which the CE1 enhancer was removed (*33*), leaving CE2 160 kb from the *Car2* promoter as the sole enhancer in the ∼400 kb TAD (**fig. S5D**). We then inserted CTCF cassettes comprising three co-oriented motifs (orientation nomenclature in **fig. S5C**), either upstream of the promoter, downstream of the enhancer, or at both locations (**Fig. 3A and B, and fig. S6**). As previously (*33*), we quantified *Car2* expression from the engineered allele (GFP) by flow cytometry, using the unmodified allele (RFP) as a control for internal normalization (see Methods).

**Figure 3.**
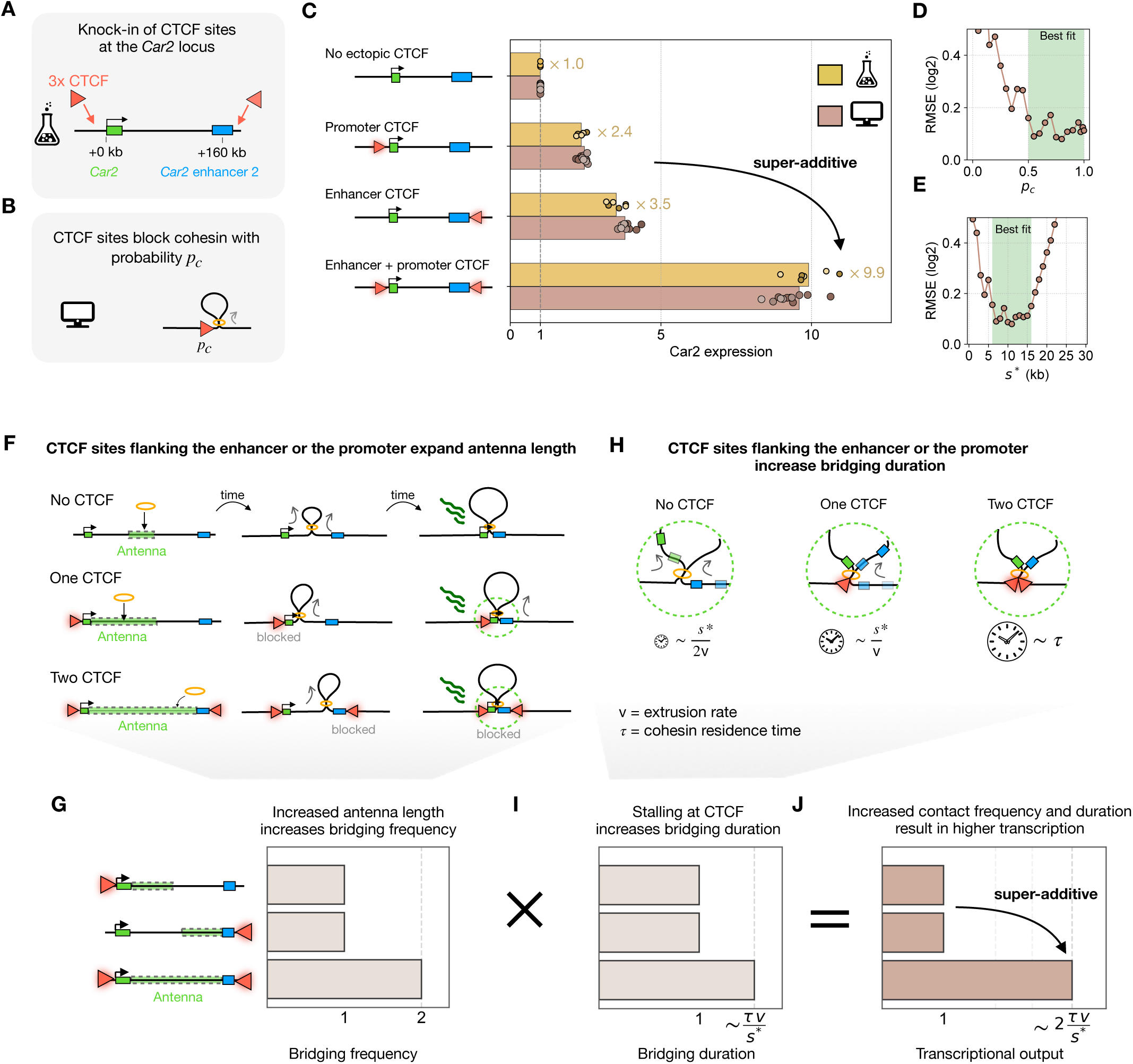
Cohesin bridging explains how flanking CTCF sites facilitate enhancer-promoter communication. **(A)** Schematic of CTCF (red arrows) site insertions near the *Car2* promoter or enhancer. CTCF arrowheads indicate the orientation of the inserted CTCF motifs. See fig. S5, fig. S6 and Methods for additional details. **(B)** In simulations, CTCF sites are represented as fixed genomic positions that capture cohesin upon passage with probability pc. **(C)** Simulations quantitatively recapitulate the facilitating effect of enhancer-and promoter-proximal CTCF sites measured experimentally, including their super-additive effect when combined. For all experiments, the normalized expression of the edited locus is shown relative to the baseline locus without ectopic CTCF sites (see Methods). Bars indicate the average across two independent clones (dark/light color) and each point is an individual flow replicate. (*n* = 3 flows from independent cultures of each clone). Predictions from the cohesin bridged model are shown using processivity 𝜆 = 120 kb, d = 150 kb. Each point corresponds to a distinct simulation with varying values of the distance threshold for activation *s\** and CTCF capture probability *pc*. For each value of *s\** within the best-fitting range (5-15 kb; see panel D), the corresponding best-fitting *pc* is shown. **(D-E)** Model fit as a function of the CTCF capture probability *pc* (D) or distance threshold *s\** (E), quantified by the root mean squared error (RMSE) between simulations and experimental data (see Methods). For each value, the best-fitting simulation is shown. The shaded region indicates the range of values yielding good agreement with the data (RMSE < 0.2) and corresponds to the parameter space used in (C). **(F)** CTCF sites flanking the enhancer and/or promoter expands the antenna length, thereby increasing bridging frequency. **(G)** CTCF sites flanking the enhancer and/or promoter increase bridging duration. With no or only one CTCF, cohesin never stalls completely, and the duration of a bridge is the time for the elements to drift past each other–which is inversely proportional to the velocity of each cohesin leg *v*. When CTCF is at both elements, the contact duration becomes the residence time of cohesin (𝜏). **(H-J)** Schematic showing that facilitating CTCF sites act additively on contact frequency through expansion of the antenna (H) and increase contact duration (I). These effects combine multiplicatively, explaining the super-additive transcriptional output in experiments and simulations (J).

Experimentally, inserting CTCF sites either ∼1.5 kb upstream of the *Car2* promoter or immediately downstream of the CE2 enhancer increased transcription by ∼3-fold, consistent with enhancer facilitation. Combining both insertions increased transcription super-additively by ∼10 fold **(Fig. 3C and fig. S6, B and C)**. We thus tested whether the cohesin-bridging model can quantitatively account for CTCF-mediated facilitation.

#### Facilitation emerges from cohesin-bridged enhancer-promoter communication

We configured our model to reproduce the genomic layout of the modified *Car2* locus by inserting extrusion barriers at the enhancer and promoter *in silico.* Each barrier represents a CTCF cassette with the blocking probability 𝑝_!_(**Fig. 3B** and Methods). Using the extrusion parameters that we constrained above (processivity 𝜆 = 110 kb, separation *d* = 150 kb), we varied the CTCF blocking probability 𝑝_!_ and the distance threshold *s\**. Not only did the model readily reproduce facilitation by either single CTCF insertion, but it also quantitatively recapitulated the super-additive effect of the double insertion of a CTCF at the enhancer and promoter (**Fig. 3C**). The best models had *s\** ∼5-15 kb and indicated that the CTCF cassette blocks cohesin with 𝑝*_c_*> 0.5 (**Fig. 3, D and E, and fig. S7**). The slight asymmetry in facilitation between promoter-and enhancer-flanking CTCF sites (2.4x and 3.5x, respectively) is explained by an endogenous CTCF site upstream of the promoter, which we previously showed contributes modestly to transcription (**fig. S5A** and (*33*)) and which we modeled as an additional weak barrier (𝑝*_c_* = 0.2; see Methods) present in all simulations.

#### Flanking CTCF facilitates distal regulation by increasing frequency and duration of cohesin bridges

The physical basis for facilitation can be understood considering the concept of antenna introduced above. Without CTCF, the antenna for symmetrically extruding cohesin is restricted to the central region of the enhancer-promoter interval. A CTCF barrier flanking the promoter and oriented toward the enhancer blocks cohesin at the promoter, allowing cohesins that load anywhere in the proximal half of the enhancer-promoter interval–not just the central region–to eventually bridge the enhancer. This expands the antenna to the entire proximal half of the enhancer-promoter interval (**Fig. 3F and fig. S8A**), thereby increasing bridging frequency. The same reasoning applies for facilitation by a CTCF barrier flanking the enhancer. With CTCF barriers at both the enhancer and the promoter, cohesin loading at any position within the enhancer-promoter interval can eventually bridge the two elements. This further doubles the length of the antenna (**Fig. 3F and fig. S8B**) and hence doubles the frequency of enhancer-promoter bridging (**Fig. 3G**).

Beyond increasing bridging frequency, facilitation by CTCF also increases bridging duration. Without CTCF barriers, the duration of an enhancer-promoter bridge is the time the elements are held together below *s\** as cohesin translocates: such bridges last ∼*s\**/2*v* (**Fig. 3H**), corresponding to ∼10 seconds, or even longer (∼100 seconds) if further translocation is blocked by collisions with other cohesins. When both promoter and enhancer are flanked by CTCF sites, both sides of cohesin will stall and the enhancer-promoter bridge lasts for the cohesin residence time 𝜏 (∼10-20 minutes) (**Fig. 3H and I**)–consistent with live-cell measurements (∼10-30 minutes) (*42, 43*).

Importantly, even a weak flanking barrier can increase transcription output, especially in the genomic context where *s\** is small: e.g. 𝑝*_c_* = 0.1, *s* =* 2 kb, *L =* 200 kb would boost transcription 2.5-fold (**fig. S8, C and D**). These considerations extend beyond CTCF to any cohesin-blocking barrier such as, possibly, active promoters (*65*)–including eRNA transcription start sites in enhancers.

Together, our analyses indicate that promoter-and enhancer-flanking CTCF barriers facilitate long-range regulation through two coupled mechanisms: by expanding the genomic region from which cohesin can productively bridge the two elements, and by prolonging the duration of their encounters. This in turn provides the physical basis for super-additivity (**Fig. 3, G and I-J**).

### Biophysical basis for transcriptional insulation by intervening CTCF barriers

We next asked whether the cohesin-bridging model can also reproduce transcriptional insulation, when CTCF sites are inserted between the enhancer and promoter (**Fig. 4A**). We knocked in two divergent 3x CTCF cassettes between the enhancer and the promoter at the *Car2* locus, resulting in more than 50-fold downregulation of *Car2* (**Fig. 4 and fig. S9, B and C**). For comparison, when inserted flanking to the promoter and enhancer, CTCF sites caused ∼10-fold upregulation (**Fig. 3**). These observations highlight how the configuration of CTCF sites around an enhancer and promoter can tune transcriptional output over more than two orders of magnitude.

**Figure 4.**
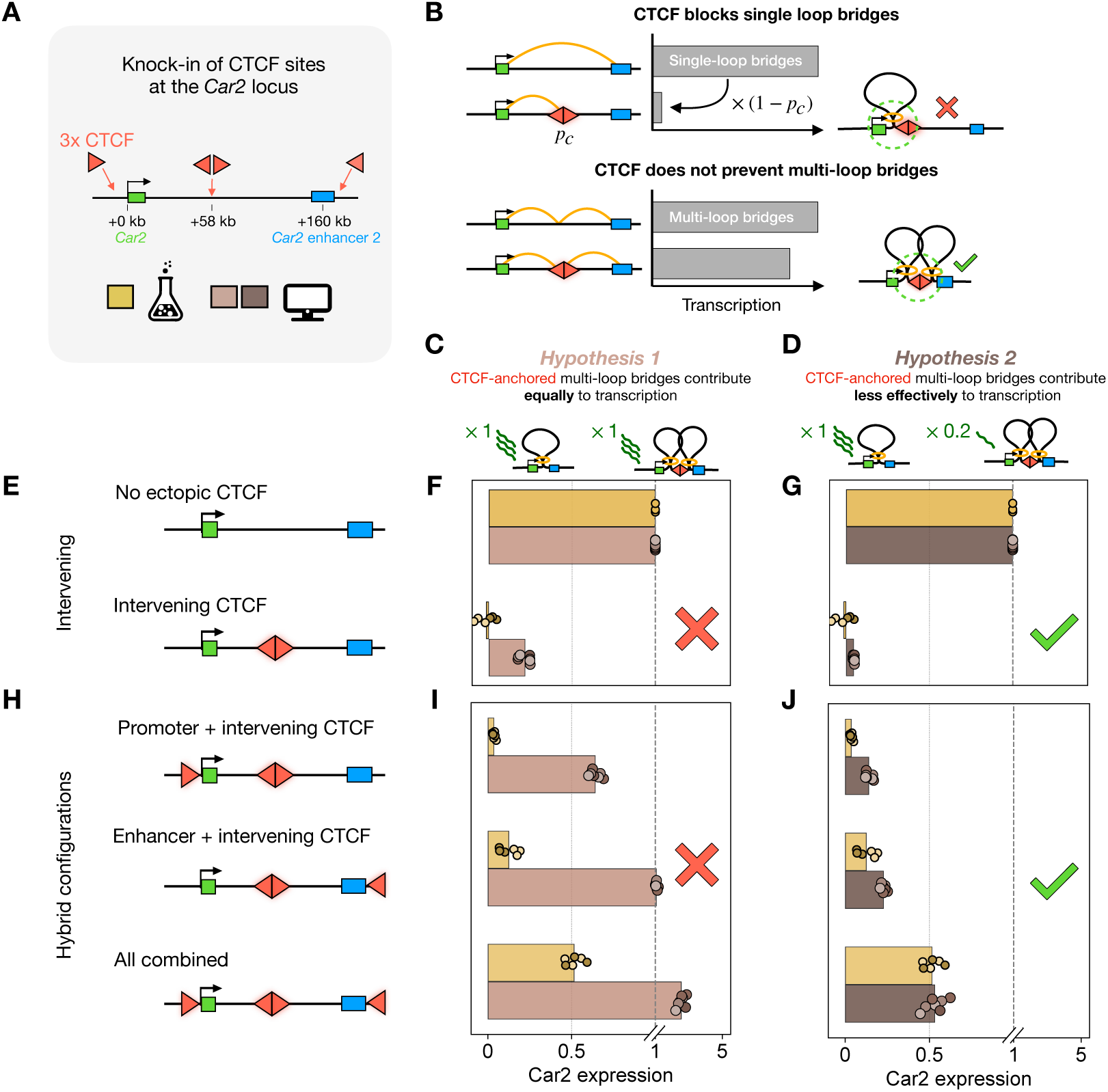
Mechanistic requirements for functional enhancer-blocking by intervening CTCF sites. **(A)** CTCF site insertions at the *Car2* locus. Arrowheads indicate the orientation of the inserted CTCF motifs. See fig. S17A and Methods for additional details. **(B)** Schematic illustrating how intervening CTCF sites block cohesin-mediated enhancer–promoter bridging. By halting extrusion, CTCF prevents a single cohesin complex from forming a loop that spans both elements, thereby strongly reducing the contribution of single-loop bridges to transcription. In contrast, multiple cohesin complexes loaded on either side of the CTCF sites can cooperate to form a CTCF-anchored rosette that span the barrier, thereby supporting enhancer-promoter bridging despite the intervening CTCF site. **(C and D)** Mechanistic hypotheses explored in the simulations. **(E)** Insulation by intervening CTCF sites. **(F)** Models assuming that all cohesin bridges contribute equally to transcriptional output underestimate transcriptional insulation by intervening CTCF sites. For experiments, the normalized expression of the edited locus is shown relative to the baseline locus without ectopic CTCF sites. Bars indicate the average across two independent clones and each point is an individual flow replicate (n = 3 flows per clone). For models, each point corresponds to a distinct simulation parameter set selected to simultaneously fit insulation and facilitation cell lines (see fig. S10), and bars indicate the mean across simulations. **(G)** Same as (F), but assuming that bridging through CTCF-anchored rosettes contributes less effectively to transcriptional output than other cohesin bridges (20%, Methods). Under this regime, the model recapitulates the strong insulation observed experimentally. **(H)** Hybrid genomic layouts combining enhancer-promoter flanking and intervening CTCF sites. **(I and J)** While models that assume all cohesin bridges contribute equally to transcription underestimate insulation in hybrid genomic layouts, models where the contribution of CTCF-anchored rosettes are reduced (to 20%) successfully recapitulate experimental observations.

To model insulation, we first considered how intervening CTCF sites affect cohesin bridging. While such barriers can efficiently block bridges formed by a single cohesin, they cannot prevent bridge formation by two or more loops stacked on opposite sides of the barrier (referred to here as CTCF-anchored “rosettes” (**Fig. 4B**), to distinguish them from multi-loop bridges formed by cohesin collisions in the absence of CTCF anchoring, see **fig. S10A**). The extent to which these rosettes support enhancer-promoter communication is therefore expected to determine how strongly intervening CTCF sites can insulate transcription (**Fig. 4C** and **D**). Two views exist on this question. Loop stacking between CTCF boundaries has been proposed to enable long-range regulatory communication (*66, 67*), implying CTCF rosettes can in some cases transmit regulatory information. Conversely, recent evidence suggests that CTCF hub formation and transcriptional regulation can be decoupled (*68*), indicating that in other cases rosette-mediated proximity is poorly conducive to enhancer-promoter activation. To distinguish between these possibilities, we simulated two scenarios: (i) all cohesin bridges contribute equally to enhancer-promoter communication, including single-loop bridges and rosettes (**Fig. 4C**); and (ii) CTCF-anchored rosettes contribute less effectively (**Fig. 4D**).

When all cohesin bridges were assumed to contribute equally to enhancer-promoter communication, simulations predicted only 5-fold down-regulation upon inserting the intervening CTCF sites (**Fig. 4, E and F**), even with a fully impermeable barrier (*p_c_* = 1), falling short of the observed >50-fold transcriptional insulation. In contrast, considering that CTCF rosettes contribute less effectively reproduced quantitatively the >50-fold transcriptional insulation observed experimentally (**Fig. 4G**; best fits were obtained after assigning a relative contribution ≤ 0.2-fold, **fig. S10, A to B**). This strong insulation further constrains model parameters, requiring high CTCF capture probabilities (*p_c_* > 0.95), which, together with constraints from facilitation, yields *s\** = 10-15 kb (**fig. S10, C to E**). Note that multi-loops arising from cohesin collisions, and not anchored at CTCF sites, are assumed to be equally efficient in mediating enhancer-promoter communication (**fig. S10A**). Altogether, these results show that cohesin bridging can explain strong insulation by CTCF sites and imply that proximity through CTCF-anchored rosettes is poorly conducive to enhancer-promoter activation in our experimental set up (**fig. S11**). Several factors could account for the reduced functional contribution of rosettes, including steric interference caused by CTCF and cohesin clusters that may hinder enhancer-promoter encounters, or increased enhancer-promoter separation due to partial nucleosome loss within CTCF arrays.

Having established that cohesin bridging unifies both the facilitating and insulating roles of CTCF, we challenged our model predictions with hybrid configurations in which these two effects combine. We engineered cell lines in which the intervening CTCF cassette is combined with promoter and/or enhancer-flanking CTCF sites (**Fig. 4H**). The cohesin bridging model (with parameters constrained within the ranges defined above) accurately reproduced all three combinatorial layouts, with best agreement achieved for a rosette contribution of ∼0.2 (**Fig. 4, H to J**). Together, these results indicate that enhancer-promoter regulation by cohesin bridging can accurately predict how complex arrangements of CTCF sites, separated by over 150 kb, can tune transcriptional output over several orders of magnitude.

### 3D contacts cannot be fully insulated by blocking loop extrusion

The accuracy of our transcriptional predictions from cohesin bridging dynamics questions the prevailing view that regulatory communication relies on general 3D proximity, irrespective of the mechanism driving enhancer-promoter encounters (*69–71*). We therefore investigated whether modulation of overall 3D proximity–in addition to cohesin bridging–can account for how cohesin and CTCF implement transcriptional insulation and enhancer facilitation. We simulated the 3D dynamics of the *Car2* locus as a chromatin fiber subject to polymer fluctuations, loop extrusion, and CTCF barriers (**Fig. 5A**). In this model, transcription is proportional to the fraction of time the enhancer and promoter are in 3D proximity, i.e. closer than a radius of contact *R_c_*. Since the spatial distance at which an enhancer can activate a promoter is unknown, we explored a broad range of *R_c_* (∼20–220 nm, spanning the size of molecular complexes up to transcriptional condensates (*72, 73*)). For each *R_c_*, we swept extrusion parameters to best match transcription for all CTCF layouts.

**Figure 5.**
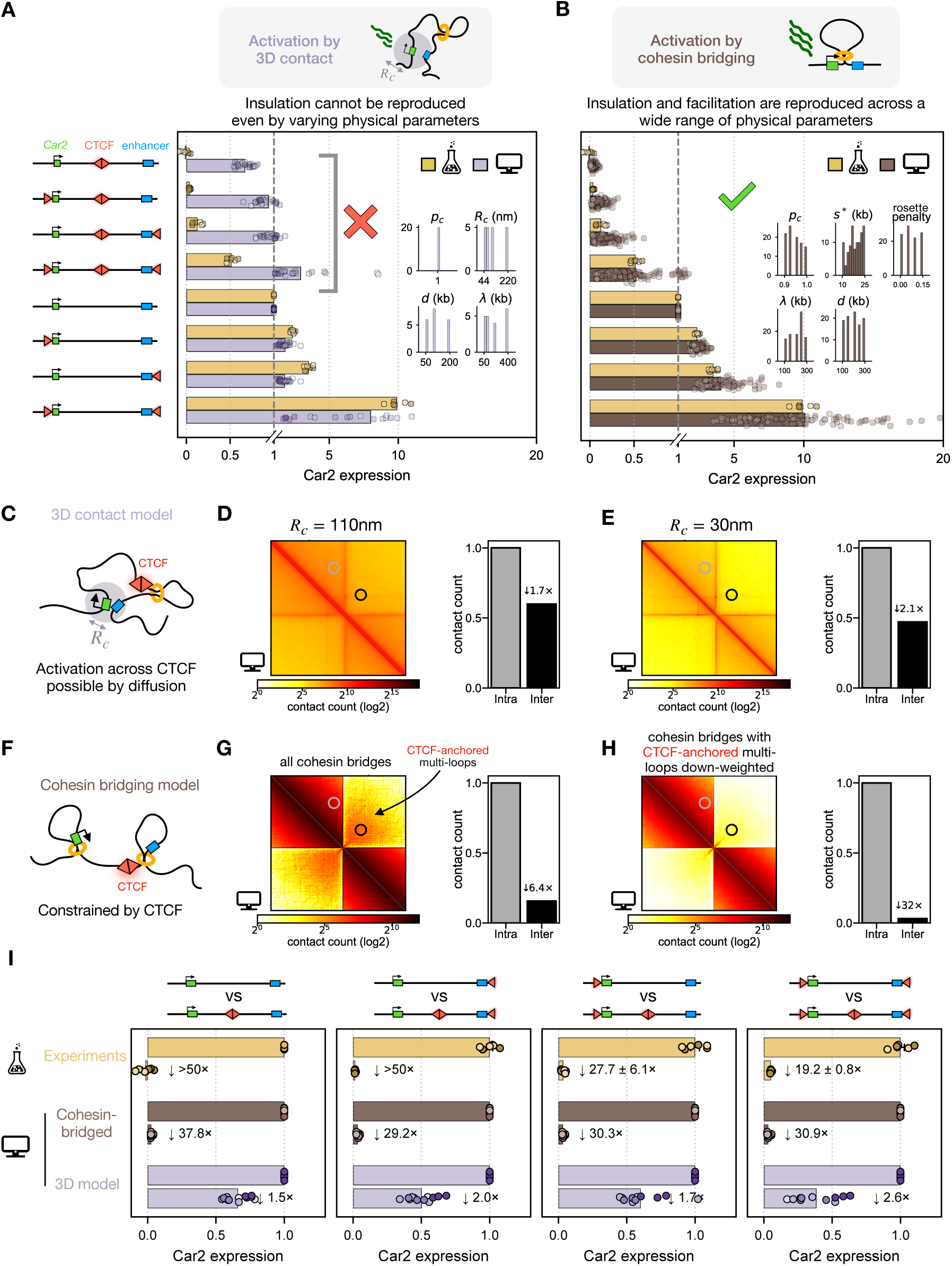
Activation via simple 3D contact fails to account for insulation, facilitation, and their combination. (A) Models predicting enhancer-promoter activation by 3D contacts failed to reproduce the strong transcriptional insulation by intervening CTCF sites measured experimentally. Dots represent individual simulations with distinct parameter combinations; bars indicate the means over these simulations. Simulations were performed over a parameter grid (inset), with the top three best-fitting simulations shown for each value of the contact radius *Rc*. The CTCF capture probability was set to *pc* = 1 to maximize insulation. Experimental data is reproduced from Fig. 3 and Fig. 4. (B) Models predicting enhancer-promoter activation by cohesin bridges quantitatively reproduce both insulation and facilitation. Dots and bars are as in (A). Simulations were sampled over a parameter range centered around the optimal fit (inset). Experimental data same as in (A). (C) Schematic illustrating that diffusion-mediated contacts in the 3D model cannot be fully prevented by intervening CTCF sites. (D) Simulated contact map from the 3D model with a large contact radius (*Rc* = 110 nm). Inter-TAD contacts are only modestly reduced relative to intra-TAD contacts, explaining the weak transcriptional insulation in models predicting enhancer-promoter activation by 3D contacts. (E) Same as (D) but for small contact radius *Rc* = 30 nm. Inter-TAD contacts remain only modestly reduced. (F) Schematic illustrating how intervening CTCF sites prevent enhancer-promoter bridging by cohesin. (G) Simulated cohesin-bridged contact map, where contacts are defined as pairs of loci bridged by cohesin loops (see Methods). While CTCF-anchored rosettes can support low but detectable inter-domain contacts, inter-TAD contacts are strongly reduced relative to intra-TAD contacts. (H) Same as (G) but the contribution of CTCF-anchored rosettes are down-weighted to 20% as in Fig. 4. This further depletes inter-vs. intra-TAD contacts (>30-fold). (I) Panels directly comparing each genomic layout with flanking CTCF sites (normalized to 1) to their counterpart with additional intervening CTCF sites–bars and dots same as in (A) and (B).

Such a model of promoter activation by 3D contact with the enhancer can reproduce facilitation by enhancer-and promoter-proximal CTCF sites (**Fig. 5A**), provided *R_c_* is small enough. However, this model fails to reproduce transcriptional insulation by intervening CTCF sites, achieving at most 2.5-fold down-regulation by a completely impermeable intervening CTCF barrier (𝑝_!_ = 1). This falls short of the >50-fold insulation observed experimentally–even for small *R_c_* (**Fig. 5A**)–indicating a fundamental limitation for this type of interaction in achieving insulation. This contrasts with the predictions obtained from modeling enhancer-promoter activation through cohesin bridging, which readily capture the transcriptional effects of CTCF sites across two orders of magnitude (**Fig. 5B**).

The reason 3D contact models underestimate transcriptional insulation becomes clear from the corresponding simulated contact maps: even a perfect extrusion barrier only reduces inter-TAD 3D contacts by at most ∼2-3 fold (**Fig. 5, C to E**), consistent with Hi-C and micro-C showing only ∼2-3 fold physical insulation across TAD boundaries genome-wide (*16, 74, 75*). This means spatial contacts arising from polymer fluctuations remain sufficiently frequent across CTCF barriers (**Fig. 5, C to E**). While spatial contacts cannot be substantially insulated by CTCF, cohesin bridges across CTCF barriers are strongly suppressed (**Fig. 5, F to H)**, explaining why a model based on bridge dynamics accurately captures the transcriptional modulation imposed by CTCF (**Fig. 5B**). This contrast is most apparent when CTCF layouts with and without intervening sites are compared directly: while the 3D contact model consistently underestimates insulation across all configurations, the cohesin-bridging model accurately reproduces experimental downregulation in each case (**Fig. 5I**).

We noticed that upon combining all three CTCF insertions, Car2 expression recovered to ∼50% of the unmodified locus (**Fig. 4, H to J**). Our model indicates that while CTCF-anchored rosettes contribute to this recovery, as previously suggested (*66*), insulator leakage also makes an important contribution: occasional cohesins can traverse the intervening CTCF sites, and flanking CTCF sites amplify this residual communication. In this context, ∼50% of the unmodified locus still corresponds to a ∼20-fold reduction relative to the genomic layout with the promoter-and enhancer-flanking CTCF sites (**Fig. 5B)**. Thus, intervening CTCF sites apply a strong suppressive factor to transcriptional output regardless of whether flanking sites are present (**Fig. 5I**).

Since only cohesin bridges–not spatial contacts–can be effectively blocked by CTCF, the potency of CTCF as a transcriptional insulator further establishes cohesin bridges, not mere spatial proximity, as the physical basis for enhancer-promoter communication at the *Car2* locus.

#### Robustness of cohesin-bridged enhancer-promoter communication

Next, we addressed whether our conclusions depend on precise parameter tuning. In addition to parameters of loop extrusion, critical parameters of the cohesin-bridging model (*s\** and *p_c_*) can be established from just two CTCF layouts: one insulating, one facilitating arrangement, with other experiments predicted without further fitting (**fig. S10F and Methods**).

To assess sensitivity more broadly, we randomly sampled parameters around their best-fitting values. Predicted transcriptional behaviors remained remarkably robust, and each set of randomized parameters correctly ranked transcriptional effects across all CTCF layouts (Spearman’s *ρ* = 1.00; **Fig. 5B and fig. S12**). This robustness indicates that model predictions reflect the underlying molecular logic by which CTCF modulates cohesin bridges, rather than being simply caused by fine-tuning. These findings also indicate that insulation and facilitation by CTCF consistently emerge from this control system, making the model applicable to other cell types or cell cycle stages. This remains true even when extrusion parameters may vary across cells due to differences in the expression of CTCF, cohesin, or its co-factors.

#### CTCF sites insulate transcription bidirectionally by preventing loop extension in two distinct ways

Our results establish that intervening CTCF sites are potent transcriptional insulators because they prevent the formation of cohesin bridges between enhancers and promoters. Yet CTCF sites are inherently polar: the DNA-bound CTCF protein blocks cohesins approaching their front side, where the CTCF N-terminus engages cohesin, halting extrusion and potentially inhibiting cohesin unloading (*42, 76–79*). This causes genome folding patterns to depend on the orientation of CTCF binding sites (*3, 4, 19, 80–83*). Whether cohesin simply passes through the backside of CTCF sites (C-terminal side of the CTCF protein), as generally assumed (*3, 4, 42, 82–86*), remains less clear. We therefore tested how different models of backside CTCF-cohesin encounters account for three experimental datasets: insertions of a single oriented CTCF motif at multiple positions across the *Car2* locus (**Fig. 6A and fig. S13**), published *Sox2* reporter measurements in which its enhancer is repositioned across a fixed directional CTCF site (*26*) (**Fig. 6B**), and published Micro-C pile-ups around isolated, uniformly oriented CTCF sites (*74*) (**Fig. 6C**).

**Figure 6.**
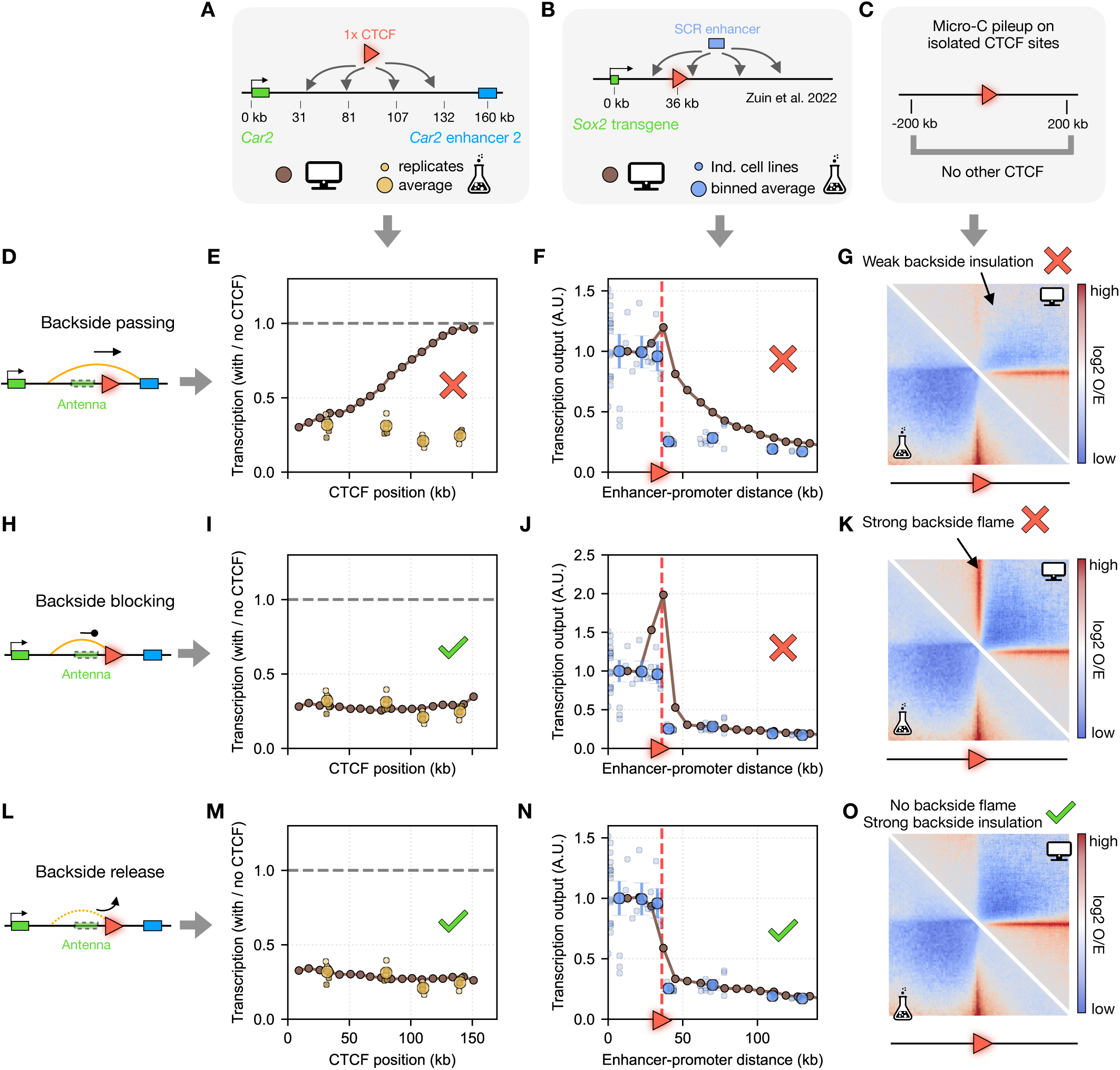
A single CTCF binding site can act as a bidirectional insulator via backside release of cohesin loops. **(A–C)** Experimental systems used to infer the outcome of cohesin encountering the backside of a directional CTCF site: (A) *Car2* expression after inserting a CTCF site at various positions across the locus, (B) *Sox2* reporter expression after enhancer relocation across a fixed CTCF site (data from (*26*)), and (C) genome-wide micro-C pileups at isolated CTCF sites in mouse ES cells to probe chromosome folding (data from (*74*)). **(D–G)** Backside passing predicts position-dependent insulation at *Car2*, a gradual rather than sharp decline in *Sox2* transcription after enhancer relocation across the CTCF site, and weak physical insulation on the CTCF backside in micro-C–inconsistent with all three experimental systems. For experiments at *Car2*, transcription is shown as the ratio of expression from the locus carrying the ectopic CTCF site to that without. **(H–K)** Backside blocking recovers position-independent insulation at *Car2* but yields spurious enhancer facilitation near the CTCF site at the *Sox2* reporter locus, and erroneously predicts symmetric micro-C stripes around CTCF sites. **(L–O)** Backside loop release correctly recapitulates all three experimental systems: position-independent insulation at *Car2*, sharp transcriptional drop of *Sox2* with enhancer insertions across the CTCF site, and symmetric micro-C insulation without a backside stripe. For experimental data in (E), (I), and (M), circles represent the average across all replicates (points) collected from 2 independent clones (n = 3 flows per clone). Best fits for backside release models used *pfrontside-capture* = *pbackside-release* = 0.7 (*Car2*) or 0.6 (*Sox2*) – reflecting the use of distinct CTCF motifs in each system.

If cohesin simply passed through the backside of CTCF sites (**Fig. 6D**), the cohesin-bridging model predicts that transcriptional insulation should depend strongly on both the position and orientation of a CTCF motif within the enhancer-promoter interval. Insulation should be strongest when the CTCF motif faces the larger segment of the interval, where it can intercept more cohesins, and weakest when most of the interval lies on the permeable backside (**fig. S14A**). Contrary to this prediction, transcriptional insulation was comparable for all single CTCF site insertions across *Car2* (**Fig. 6E and fig. 14B)**. Backside passing was equally inconsistent with data of the *Sox2* reporter, where it predicts a gradual decline in transcription as the enhancer is relocated across the CTCF site, while experiments exhibited a sharp drop (**Fig. 6F and fig. S15A**). Backside passing also failed to reproduce the sharp, symmetric insulation pattern in Micro-C pile-ups, predicting weaker physical insulation than what is observed (**Fig. 6G**). Any viable model must therefore prevent cohesin from crossing CTCF in either orientation.

Bidirectional blocking, where CTCF captures cohesin from both front-and backside, would fulfill this requirement (**Fig. 6H and fig. S14C**). While consistent with the uniform transcriptional insulation across the *Car2* locus (**Fig. 6I and fig. S14D**), bidirectional blocking however predicts that backside encounters would also stall cohesin and stabilize loops, causing facilitation when the enhancer is close to the CTCF site–which is inconsistent with the *Sox2* reporter data (**Fig. 6J and fig. S15B**). Moreover, bidirectional blocking would create symmetric stripes on both sides of CTCF sites in contact maps, in direct contradiction with Micro-C experiments (**Fig. 6K**).

To satisfy all three experimental constraints, the backside must prevent cohesin passing without anchoring it. These requirements would be satisfied if, instead of blocking loop extension, backside encounters caused release of the DNA loop–for example through cohesin unloading or disengagement of one DNA fiber (**Fig. 6L**). In this backside release model, a directional CTCF motif would prevent cohesin passage in either orientation, but with distinct outcomes: frontside encounters block and retain the loop, while backside encounters release it (for orientation nomenclature of CTCF sites see **fig. S5C**). Since neither orientation allows cohesin to pass, this model correctly predicts position-independent insulation at *Car2* (**Fig. 6M and fig. S13E**), the sharp drop in *Sox2* transcription across a fixed CTCF site (**Fig. 6N and fig. S15C**), and the asymmetric stripe pattern with symmetric insulation edges in Micro-C (**Fig. 6O**). Frontside blocking and backside release thus both implement transcriptional insulation while leaving different imprints on genome folding. It remains to be investigated if the probabilities of backside release and frontside blocking can vary across CTCF motifs and to what extent they can vary independently of each other across genomic contexts. Beyond establishing that frontside blocking and backside release are distinct outcomes of CTCF-cohesin encounters, this work illustrates how the cohesin-bridging framework can reveal unexpected mechanistic principles of chromosome organization and long-range transcriptional control.

## Discussion

Altogether, our work puts forward the idea that not all instances of 3D chromosomal proximity are functionally equivalent for enhancer–promoter communication. Instead, transcription relies on a subset of contacts in which cohesin transiently bridges distal regulatory elements. This view addresses a long-standing challenge in interpreting chromosome conformation data from Hi-C–family methods, by establishing that most detected contacts do not transmit regulatory information. Our quantitative framework unifies several key features of long-range transcriptional regulation, including the exponential decay of enhancer activity with genomic distance (*26*), strong insulation by intervening CTCF sites despite modest changes in 3D contact frequencies (*16*), and facilitation by CTCF sites flanking enhancers or promoters (*24, 37, 82*).

What makes cohesin bridges more effective than diffusion-driven 3D contacts for long-range enhancer–promoter communication? First, by reducing the effective distance between the enhancer and the promoter, cohesin bridging confines the enhancer and promoter to a smaller search volume, thereby increasing their local concentration and contact probability. Using the scaling of contact probability with genomic distance (*P(s) ∼s^-1^*), a nearly 10-fold reduction (from *L* to *s\**) translates into ∼10-fold increase in contact probability. In this sense, cohesin bridging acts similarly to entropic or template catalysis in chemistry. Second, cohesin bridging extends the duration of enhancer–promoter encounters. This extension of encounter duration, in turn, can be regulated by cohesin cofactors that control extrusion dynamics (residence time, velocity) (*39, 87–89*) and by interactions with boundaries, such as CTCF (*21*) or active promoters (*65*). For *s\** ∼10–20 kb, corresponding to a spatial distance of ∼80–100 nm, one expects bridges to last ∼10–100s. Consistent with this estimate, direct live-cell microscopy observations of cohesin bridging events was achieved in an independent study, which reached highly concordant conclusions regarding their role in enhancer regulation (*41*). These bridges can last up to tens of minutes when stabilized by CTCF (*42–44*), far exceeding the transient contacts expected from random 3D polymer collisions (*41*)–though the exact duration of such collisions remains to be precisely determined given the strongly subdiffusive nature of chromatin motion (*90*). Together, the increased contact probability and extended encounter duration make cohesin bridges especially potent in bolstering enhancer-mediated transcription.

What is the meaning and molecular origin of the genomic separation threshold *s\**? Our estimate of *s\** ≲20 kb at the *Car2* locus is remarkably congruent with the distance where enhancer action stops relying on cohesin extrusion at this locus (*33*). We anticipate the exact value of *s\** may depend on genomic and chromatin context, potentially making some loci more or less dependent on cohesin and CTCF.

While our experiments focused on enhancer action across distances comparable to cohesin processivity (∼100–200 kb), cohesin can also support enhancer activity over several hundreds of kilobases (*36, 91*). Multi-loops formed by collided cohesins can bridge distal elements (**fig. S3 and fig. S11**), but such interactions are relatively infrequent and short-lived. Consequently, facilitation by enhancer-and/or promoter-proximal CTCF sites is likely essential for cohesin extrusion-based regulatory communication beyond several hundreds of kilobases (*21, 39, 64, 92*).

Our analyses revealed a previously unrecognized outcome of cohesin-CTCF encounters: release following a C-terminal encounter, rather than passing as previously assumed (*3, 4*). Since both frontside blocking and backside release prevent cohesin from crossing the CTCF site, both orientations can implement transcriptional insulation–yet with distinct effects on Hi-C maps. In contrast, facilitation by promoter-or enhancer-flanking CTCF sites relies on cohesin stopping and retention, and therefore only occurs upon frontside encounters, where the N-terminus of CTCF interacts with cohesin to stabilize it (*76, 78*). Backside release remains consistent with the observation that Hi-C dots preferentially engage convergent pairs of CTCF sites (*4, 80, 82, 83*), as these correspond to frontside encounters (*76–78*). The probability of backside release need not equal that of frontside blocking and both are expected to vary across CTCF motifs and genomic context (*93, 94*)–determining the extent to which a given CTCF site can block, permit or bolster regulatory communication.

Backside release explains why N-terminal mutations of CTCF, which impair frontside blocking and abolish Hi-C dots and stripes, do not fully disable TAD insulation (*76, 78*). Our work opens the door to investigating the structural basis for backside release and how it may vary across CTCF motifs and genomic contexts. *In vitro* studies did not report loop release upon backside encounters (*95, 96*), possibly due to the lack of necessary co-factors in the reconstitution. The CTCF C-terminus is unlikely to be involved as its removal in cells does not prevent CTCF from insulating TADs or from retaining cohesin (*76, 77, 97*). Finally, since the enhancer-blocking effect of CTCF sites can depend on orientation in other reporter assays (*31, 98*), we anticipate that motif composition or genomic context may tune, and possibly decouple, the probabilities of frontside blocking and of backside release.

While our work employed CTCF site insertions to disrupt cohesin trajectories, we anticipate other types of extrusion barriers can also modulate enhancer-promoter bridging. Transcription itself can act as a moving barrier to cohesin (*65, 99, 100*) and intervening promoters can act as insulators (*101–103*). Other transcription factors can cause cohesin retention outside CTCF sites (*104, 105*), a sign of cohesin blocking (*93*), including at some developmental enhancers (*106–109*). This raises the intriguing possibility that in some specialized contexts transcription factor binding might couple local commissioning of enhancers with long-range regulatory communication by simultaneously creating a cohesin barrier–even weak.

Even though cohesin bridging is only one of several processes contributing to long-range enhancer-promoter communication (*59, 60*), the concept of *s\** helps explain how alternative mechanisms can operate. First, when an enhancer and promoter are proximal (separated by less than *s\**), their communication becomes largely cohesin-independent (*33, 36, 37*). Second, tethering through “sticky” factors, such as transcription (co)-activators or adaptor proteins (*59, 60*), binding within *s\** of either the enhancer or promoter would keep them in proximity after their first encounter–whether it was extrusion-or diffusion-initiated (*38, 41, 58, 110–114*). Such mechanisms can buffer some loci against perturbations to loop extrusion (*33*). Our model does not exclude additional transcriptional roles for CTCF and cohesin beyond enhancer-promoter bridging (*34, 115–117*). Another role of cohesin, and broadly loop extrusion, can be to facilitate encounters between distal and slowly diffusing genomic loci (*90, 118, 119*), as recently demonstrated in yeast (*120, 121*). Yet, as we demonstrate, cohesin-bridged interactions remain uniquely sensitive to CTCF insulation–creating the unique opportunity to partition regulatory landscapes that is essential for precise developmental patterning.

The cohesin-bridging mechanism could explain a broad range of regulatory phenomena (**fig. S16**). For example, facilitation of cohesin bridges by CTCF can enable targeting of genomic communication to a specific promoter while skipping intervening ones–a well-known but poorly understood feature of enhancer–promoter communication. Targeted activation of transcription can be highly precise, with a facilitating CTCF enabling the preferential selection of a single promoter within a dense cluster (with relevance for *Hox* or protocadherin clusters for example (*122*)). While these complex loci undoubtedly involve additional regulatory layers, the precision afforded by cohesin bridging may nonetheless provide unique regulatory opportunities essential to their function. As another example, the model predicts that inverting a facilitating CTCF site reduces transcription, since facilitation requires cohesin retention by the frontside of CTCF motif, which is no longer supported after inversion because of backside release (**fig. S16**).

By building from a locus with minimal regulatory complexity, our study establishes a physical basis for core aspects of long-range enhancer-promoter communication. Extending this to genome-wide predictive mechanistic models will require meeting the challenges of mapping enhancer dependencies at scale, formalizing the quantitative logic governing interactions among multiple regulatory elements, and integrating the full diversity of cohesin-dependent and-independent long-range regulatory mechanisms (*59, 123*).

## Material and methods

### Experiments

#### Choice of CTCF sites for insertion at *Car2*

For all CTCF site insertions at the *Car2* locus, the core motif plus 10-20 bp of flanking endogenous sequence were chosen from the mouse genome. In the 3X CTCF site cassettes, individual CTCF sites were selected based on high occupancy by ChIP-seq from (*93*) and were found in TAD boundaries. The divergent cassette inserted in the intervening region between *Car2* and its enhancer additionally contained 1 kb of neutral spacer sequence from the human genome separating the 3X reverse and 3X forward oriented motifs. The 3X reverse oriented cassette inserted at *Car2* enhancer and intervening position were identical, while the forward oriented sites were all unique (**fig. S17A**). For the insertions of a single CTCF site at various positions across the *Car2* locus, the same sequence was used for each location (**fig. S17B**).

#### Plasmid construction

Plasmids were assembled using Gibson assembly (NEB HiFi DNA Assembly Master Mix E2621) or site-directed mutagenesis (NEB PNK T4 Polynucleotide Kinase M0201S, NEB Quick Ligase M2200). The list of plasmids generated in this study can be found in Supplementary Table S1 and their annotated sequence maps as Supplementary Data S1.

#### Cell culture

Parental WT E14Tg2a (karyotype 19, XY, 129/Ola isogenic background) mouse embryonic stem cells (ESCs) and subclones were cultured in 2iSL medium [DMEM + GlutaMAX + sodium pyruvate (ThermoFisher 10569044) supplemented with 15% fetal bovine serum (Gibco SKU A5256701), 550 µM 2-mercaptoethanol (ThermoFisher 21985-023), 1X non-essential amino acids (ThermoFisher 11140-050), 10^4^ U/mL of Leukemia Inhibitory Factor (Millipore ESG1107), 1 µM PD0325901 (Apex Bio/Fischer A3013-25), and 3 µM CHIR99021 (Apex Bio/Fischer A3011-100)]. Cells were maintained at a density of 0.2-1.5 x 10^5^ cells/cm^2^ by passaging using TrypLE (ThermoFisher 12605010) every 24-48 hours on 0.1% gelatin-coated dishes (Sigma-Aldrich G1890-100G in 1X PBS) at 37 °C and 5% CO_2_. The medium was changed daily when cells were not passaged. Cells were checked for mycoplasma infection every 6 months and tested negative.

#### Genome engineering

For transfection, plasmids were prepared using the Nucleobond Midi kit (Macherey Nagel 740410.5) followed by isopropanol precipitation. Knock-in constructs were not linearized. All transfections were done using the Neon system (Thermo Fisher) with a 100 µL tip and 1 million cells at 1400 V, 10 ms, 3 pulses. To create knock-in cells, ESCs were co-transfected with 3 µg of Cas9-sgRNA expressing vector and 14 µg of targeting construct. Homology arms of targeting vectors ranged from 800 base pairs (bp) - 1 kilobase (kb) in length.

After electroporation, cells were seeded in a 9 cm^2^ well and left to recover for 48 hours. Cells were plated at limiting dilution in 10 cm dishes and grown for around 8 days changing medium every other day until single colonies could be picked. Individual colonies were genotyped by polymerase chain reaction (PCR) and validated with Sanger sequencing. Homozygous or heterozygous clones were identified, expanded, validated by flow cytometry, and cryopreserved. For serial editing, resulting validated clonal lines were used to repeat the transfection process with another targeting construct.

Because the targeting constructs did not contain a resistance cassette we used sgRNAs cloned into the Cas9-2A-puro vector (pEN243, identical to pX459 Addgene 62988) and included puromycin for 48 hours in the culture medium starting 24 hours after transfection. This was not strictly necessary as transfection efficiencies were typically above 80% but it can help remove untransfected cells to increase the chance of picking edited clones.

#### Genotyping small insertions at the *Car2* locus

To genotype cell lines where the knock-in was small (<500 bp), a unique restriction site was included in the cassette. A PCR was set up using primers that were anchored outside of one homology arm and spanned the knock-in location **(fig. S17C)**. This PCR reaction was immediately digested based on the unique restriction site to distinguish between wild-type, heterozygous, and homozygous knock-ins **(fig. S17D).**

The list of cell lines, genotypes, order of transfection, and corresponding vectors is provided in Supplementary Table S1.

#### Flow cytometry

ESCs were dissociated with TrypLE, resuspended in culture medium, spun, and resuspended in 10% FBS-PBS before live cell flow cytometry on an Attune NxT instrument (Thermo Fisher). Analyses were performed using the FlowJo software. Briefly, live cells were gated using FSC-H x SSC-A followed by the isolation of singlets using FSC-H x FSC-W before extracting statistics for approximately 50,000 final gated cells. It was necessary to account for any effects in *trans* in the dual fluorescent reporter cell lines, for example, changes in *Car2* reporter expression due to variable cell density (*33*). For this, the population median expression of the edited allele was divided by the population median of the unedited allele with E14 untagged autofluorescence removed, all relative to a control cell line included in every flow. Replicate data points were obtained by growing and flowing the same cell line on different days. For each cell line, two independent clones were analyzed and are presented.

#### Analytical theory for bridging frequency by non-interacting cohesin

To interpret the distance dependence of enhancer–promoter communication, we derived an analytical expression for the bridging frequency under the assumption of non-interacting cohesins (Supplementary Text, Eq. 18). The formula was used to generate the theoretical curves shown in Fig. 2A. A detailed derivation and discussion of the underlying assumptions are provided in the Supplementary Text. Note that similar analytical frameworks have been explored previously (*124–126*) albeit in different systems and for distinct observables.

### Simulations

#### Modelling cohesin translocation dynamics on DNA

To simulate the dynamics of cohesin bridges between two distal elements, we developed a Cython-based loop-extrusion simulator, available at https://github.com/mirnylab/lefs-cython. Here we briefly summarize its logic and describe how it was used in this work.

As in previous studies (*3, 126*), chromatin is represented as a one-dimensional lattice of N sites, each site representing 1kb of chromatin, and a fixed number of cohesins N_c_ are simulated on this lattice. Each cohesin is represented by two legs, each occupying a single lattice site and representing the anchoring points of cohesin on DNA. The positions and states of cohesin legs are updated in discrete time according to the following rules:

● **Leg activity**: Each cohesin leg can be either active or inactive. At each time step, only active legs are eligible to translocate. In the simulations used here, we employed a one-sided extrusion scheme in which only one leg of a cohesin is active at a time, while the other is inactive (move_policy = 1). The identity of the active leg switches stochastically with probability k_switch_ per update step. If one leg is captured at a barrier, the opposite leg becomes and remains the active leg.
● **Translocation of the active leg**: Loop extrusion occurs by translocation of active legs away from one another along the lattice. At each step, an active leg advances by one lattice site provided that the destination site is not occupied by another cohesin, or isn’t captured by a barrier (see paragraphs about cohesin-cohesin interactions and barriers).
● **Unbinding and rebinding**: Cohesins stochastically unbind with a constant probability per time step 𝑘_&_, corresponding to a finite residence time on chromatin 𝜏 = 1/𝑘*_u_*. Upon unbinding, cohesins are immediately reloaded at new positions drawn at random from the available lattice sites.
● **Cohesin-cohesin interactions**: Cohesins obey steric exclusion, such that no two legs can occupy the same lattice site. When an active leg attempts to move to an occupied site (i.e., a site already occupied by another cohesin leg), the move is aborted and the leg is marked as *stalled*. Stalled legs remain in place and can attempt to move again in subsequent time steps. There is no bypass or displacement of other cohesins. Importantly, stalling of one leg does not prevent continued extrusion by the opposite leg, provided that the latter is active and its target site is unoccupied.
● **Extrusion barriers**: Specific lattice sites can be assigned as extrusion barriers that interact with cohesin legs in a probabilistic and direction-dependent manner. Upon encounter with a barrier, a cohesin leg can either be captured with probability p_c,f_ when approached from the frontside and p_c,b_ when approached from the backside, or induce cohesin unbinding with probabilities p_u,f_ and p_u,b_, respectively. If neither capture nor unbinding occurs, the cohesin leg passes through the barrier, provided the target site is not occupied by another cohesin (often a cohesin captured at this site). Captured legs remain immobilized, while the opposite leg may continue to extrude if unobstructed. In this implementation, once captured, the cohesin leg is not released until cohesin unbinding. The cohesin unbinding rate is not altered by CTCF capture; therefore, the lifetime of CTCF-bound cohesin is the same as that of unbound cohesin, and a loop anchored between two convergent CTCF sites has mean lifetime τ.

#### Simulation units and physical parameters

In the main text, we parameterize the system using the standard physical quantities: the cohesin extrusion velocity v, the average genomic separation between cohesins d, and the cohesin processivity λ, defined as the mean genomic distance extruded before dissociation for an unobstructed cohesin.

These physical parameters were mapped onto the discrete simulation framework as follows. Extrusion proceeds at one lattice site per update step, such that the extrusion velocity is given by 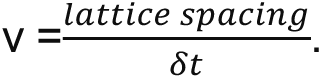 Since the lattice spacing is fixed to 1 kb, specifying v directly sets the duration of one simulation time step as 𝛿𝑡 = 1/𝑣. This mapping was used to express inter-bridging times and bridging durations in real time units when comparing to experimental measurements.

Cohesin unbinding probability k_u_ was set such that the mean genomic distance extruded before dissociation matches the specified processivity λ. In the one-sided extrusion regime used here, this corresponds to setting the unbinding probability per step to k_u_=1/λ. The average genomic separation between cohesins, d, was controlled by fixing the total number of cohesins on the lattice to N_c_ = ⌊N/d⌋.

#### Modelling enhancer-promoter bridging by cohesin and subsequent promoter activation

The enhancer and promoter were represented as contiguous intervals of length l_e_ and l_p_, separated by a genomic distance *L* (defined as the distance from the center of the promoter to the center of the enhancer). At each time step, the effective genomic separation between enhancer and promoter, s_eff_, was computed as the shortest path between the enhancer and promoter regions along chromatin, where cohesin loops can be used as shortcuts. To this end, the current configuration of cohesin loops was represented as a graph in which nodes correspond to cohesin anchor positions as well as all lattice sites within the enhancer and promoter intervals, and edges connect neighboring genomic positions as well as pairs of anchors belonging to the same cohesin. s_eff_ was then obtained as the minimal path length in this graph, computed using Dijkstra’s algorithm. Adjacent cohesin anchors were assigned zero genomic distance between them when constructing the graph. Thus, a chain of touching loops did not accrue a distance penalty of one lattice site at each junction. In addition to the effective distance, the number of cohesin loops used along the shortest path was recorded, allowing classification of bridging events as single-loop (paths involving a single cohesin loop) or multi-loop (paths involving two or more loops).

We defined a cohesin-bridge event as a contiguous time interval during which 𝑠*_eff_* ≤ 𝑠 ∗. We recorded the frequency (inverse time between the onset of two events) and duration of these events to characterize their dynamics.

Transcriptional output was assumed to be proportional to the fraction of time the enhancer and promoter spend in the bridged state, 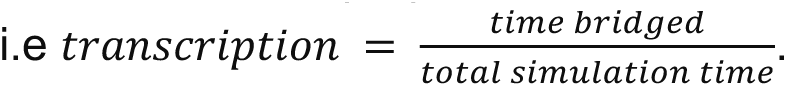.

#### Simulating transcription output function of enhancer-promoter distance and fitting to experiments at the Sox2 locus

To quantify the dependence of transcriptional output on enhancer–promoter separation, we simulated isolated E–P pairs at varying genomic distances L. For each value of L, we computed the transcriptional output as described above. We compared these predictions with measurements in which a fixed Sox2 promoter drives a reporter expression while the Sox2 control region (SCR) enhancer is randomly reinserted at different genomic positions in a neutral TAD in mouse embryonic stem cells. Each insertion generates an independent cell line, allowing transcriptional output to be measured as a function of genomic enhancer–promoter distance. We restricted our analysis to the ΔΔCTCF background, in which all CTCF sites within the TAD have been deleted. Only insertions at positive genomic distances were considered, and transcriptional output was grouped into nine fixed distance bins spanning 10–250 kb, with bin edges at 0, 27.8, 55.6, 83.3, 111.1, 138.9, 166.7, 194.5, 222.2, and 250.0 kb. Each bin was represented by its midpoint (13, 41, 69, 96, 124, 152, 179, 235, 262), and the transcriptional output assigned to that bin was taken as the mean across all insertions falling within it. Error bars were estimated as 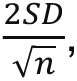 where n is the number of insertions in the bin.

We performed simulations over a broad parameter range of parameters (λ = 50 - 300kb, d = 50 - 300kb, s*=1-30kb), setting the enhancer-promoter distance at the experimental distance bins. Simulated and binned experimental transcriptional outputs were compared by fitting, for each parameter set, a single multiplicative normalization constant. This constant was obtained by minimizing the inverse-variance weighted root-mean-square error (RMSE),

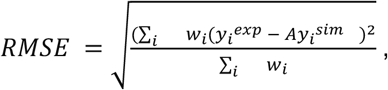

where A is a fitted multiplicative normalization constant, 𝑦*_i_^exp^* the experimental transcriptional output at bin i, 𝑦*_i_^sim^* the corresponding simulated transcriptional output and 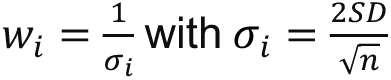 is the estimated error described above.

This normalization accounts for the arbitrary units of the experiments and allows comparison of the distance-dependent decay profiles between model and experiment. The RMSE was computed across all bins except the shortest-distance bin (13 kb). This bin was excluded because it lies near the short-distance regime, where enhancer–promoter interactions are largely independent of cohesin-mediated bridging. By restricting the fit to distances ≥ 41 kb, we focus on the regime in which transcription becomes cohesin-dependent(*33, 36, 40*).

To test whether the distance-dependent decay inferred from the Sox2-SCR dataset generalizes to other loci, we compared the best-fitting simulation curves to independent measurements at the *Car2* locus(*33*). As transcription is reported in arbitrary units that differ between systems, we fitted a multiplicative normalization constant (independently for CE1 and CE2) to rescale the experimental data. This normalization was obtained by minimizing the inverse-variance weighted RMSE between the experiments and the best fitting simulation (at d= 150kb). The rescaled data were then plotted together with the simulations to enable direct comparison of the decay profiles.

All simulations were performed on lattices of size N = 2000kb, with the enhancer placed at N/2 and promoter at N/2+L. Enhancer and promoter lengths were set to 𝑙_*_= 5 kb – reflecting the length of the H3K27ac domain at the SCR region – and 𝑙, = 0 kb. For each parameter set, simulations were run for T = 30 × 10^6 steps, after an initial burn-in period of 1000 steps to reach steady state.

#### Simulation of CTCF insertion at the Car2 locus

To model CTCF-mediated facilitation and insulation by CTCF sites at the *Car2* locus, we simulated a one-dimensional chromatin lattice of N=2000 sites, with one lattice site corresponding to 1 kb. The promoter was placed at N/2, and the remaining distal enhancer CE2 was positioned 160 kb downstream, reflecting the engineered configuration in which CE1 was deleted. Simulations were performed with one-sided loop extrusion with stochastic switching of the active cohesin leg (move_policy=1, k_switch_ =0.1, i.e one switch per 10 kb on average), enhancer and promoter lengths l_e_=3 kb, reflecting the length of the H3K27ac region at CE2, and l_p_=0 kb, cohesin separation d=150kb, processivity lambda = 110kb (fixed from the previous fit).

Engineered 3×CTCF cassettes at the promoter and enhancer were represented in silico as single barrier sites, as described above. Both inserted sites were assigned the same cohesin capture probability, p_c_ (upstream capture at the promoter, downstream capture at the enhancer). The intervening insulating cassette was modeled as two adjacent barriers with opposite orientations, thereby creating a divergent configuration that captures from both sides with the same capture probability p_c_. Constructs containing one or two intervening cassettes, as well as hybrid configurations combining flanking and intervening barriers, were generated by combining these elements on the same lattice. In all simulated cell lines, an additional promoter-proximal CTCF site was included one lattice site upstream of the promoter, with a forward capture probability fixed to 0.2. This models an endogenous CTCF site at the promoter present in all insertion experiments. In previous work, deletion of this site in a background containing both CE1 and CE2 resulted in a 20% reduction in transcriptional output(*33*). Including this CTCF captured the slight asymmetry between enhancer-and promoter-flanking CTCF-mediated facilitation (3.5x and 2.4x, respectively).

To distinguish bridging events involving intervening CTCF sites from other multi-loop configurations, we defined masks spanning the regions surrounding the intervening cassettes. These masks were used to annotate shortest enhancer–promoter paths that traversed loops anchored at the intervening CTCF sites, allowing such paths to be identified as CTCF-anchored rosettes in subsequent analyses. At CTCF sites, cohesin loading was disallowed by setting the local loading probability to zero.

All simulations included an initial burn-in period of 1000 update steps before measurements were recorded. Simulated transcriptional output was measured as the fraction of time during which the effective enhancer–promoter distance s_eff_ fell below the activation threshold s*, as described above. When intervening CTCF sites were present, configurations involving CTCF-anchored rosettes were down-weighted. Specifically, transcriptional output was defined as:

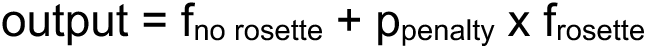

where f_no rosette_ and f_rosette_ denote the fraction of time spent in configurations with s_eff_ < s^* without and with a CTCF-anchored rosette, respectively, and p_penalty_ < 1 is a scaling factor that reduces the contribution of rosette-mediated contacts.

To reduce computational cost, path enumeration was restricted to configurations involving at most four loops.

#### Parameter fitting

To constrain model parameters, we performed a systematic exploration of parameter space by sweeping over the cohesin capture probability at inserted CTCF sites (p_c_ = 0 to 1), the activation threshold, (s* = 1 to 30kb), and the rosette penalty (p_penalty_ = 0 to 1) and compared to experimental measurements using log_2_-root-mean-square error (RMSE).

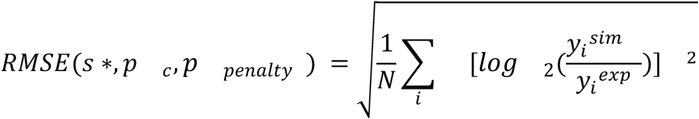

where 𝑦*_e_^exp^* is the experimental transcriptional output for cell line i, 𝑦*_i_^sim^* the corresponding simulated transcriptional output. Both simulated and experimental values were normalized by the transcriptional output of the baseline configuration lacking inserted CTCF sites, allowing direct comparison of fold changes across conditions. For the insulating configuration (CTCF at 58 kb upstream the promoter, I58) cell lines, the mean measured transcriptional output was negative due to experimental noise (one clone exhibiting fluorescence below background). Because log-ratios are undefined for non-positive values, we introduced a small pseudocount equal to the noise floor (∼0.02) for this condition when computing the RMSE.

We first constrained parameters on the facilitating cell lines i.e CTCF at enhancer, promoter, or both. Allowing for a maximum of 0.2 RMSE, we found a family of (s*, p_c_) values that fit the experimental data. However, facilitation cell lines alone do not uniquely constrain parameters.

#### Estimation of rosette penalty and CTCF capture probability

We estimated the p_penalty_ and p_c_ using the I58 cell lines. For each value of p_penalty_, we computed the RMSE between simulated and experimental transcriptional output, minimizing over all other parameters (s*, p_c_). Values of p_penalty_ < 0.2 and p_c_ > 0.95 were required to reproduce the experimentally observed level of insulation.

#### Joint constraint and parameter selection

To identify parameter sets consistent with both facilitation and insulation, we started by selecting simulation p_penalty_<0.2 to reproduce insulation of the I58 cell line. Then within these simulation we further selected (s*, p_c_) parameters whose simulated transcription had RMSE<0.2 for facilitation and RMSE<0.4 for insulation, independently to avoid over-weighting any individual cell line. These parameters were used to predict transcriptional output in hybrid configurations combining facilitating and insulating CTCF sites (E+I58, P+I58, EP+I58). Among the admissible parameter sets, the best agreement with the data was obtained for p_penalty_ = 0.2.

#### Parameter fitting on only two cell lines

We additionally performed parameter fitting using a reduced set of conditions comprising a single facilitating configuration (CTCF sites flanking both the enhancer and promoter, EP) and a single insulating configuration (I58), and assuming p_penalty_=0.2. Using the same fitting procedure as above, we inferred parameters from these two conditions and used them to predict transcriptional output across all remaining configurations.

#### Testing different CTCF-back side interaction against single CTCF insertions

To investigate cohesin–CTCF-backside interactions, we implemented three alternative hypotheses in our model and compared their predictions to experiments in which a single CTCF site was inserted between Car2 and CE2. A passing model, in which the inserted CTCF blocks cohesin only upon frontside approach, with capture probability p_c,f_=0.7, while cohesin approaching from the back side passes through. Second, a blocking model, in which the inserted CTCF blocks cohesin from both frontside and backside approaches with equal capture probabilities, p_c,f_ = p_c,b_ = 0.7. Third, a releasing model, in which the inserted CTCF blocks cohesin upon frontside approach with probability p_c,f_=0.7, but triggers unloading upon backside approach with probability p_u,b_=0.7.

The lower capture and unloading probabilities used here, relative to previous simulations, reflect the fact that these experiments involved insertion of a single CTCF motif rather than the 3× CTCF cassettes considered above; these values were also those that provided the best fit to the data. We then simulated these models using the previously constrained parameters λ = 110 kb, d=150 kb, p_penalty_=0.1, and s*=10 kb, and compared the results across the four experimentally tested insertion positions: 31, 81, 107, and 140 kb.

#### Testing different CTCF-back side interaction against enhancer insertion with intervening CTCF

To further test these hypotheses, we compared them to an independent dataset from Zuin et al. 2022, where an enhancer is inserted at various distance from a Sox2 transgene promoter, focusing on the condition in which one of the two endogenous CTCF sites is retained. In this setup, a single CTCF site is fixed, and the enhancer is repositioned at varying genomic distances. Only insertions at positive genomic distances were considered, and transcriptional output was grouped into non-uniform distance bins spanning 0–250 kb, with bin edges at 0, 15, 30, 36, 45, 60, 80, 100, 120, 140, 160, 180, 200, and 250 kb, chosen to avoid overlap with the CTCF site at 36 kb. Within each bin, the mean transcriptional output and associated uncertainty (±2 SEM) were computed, and values were normalized to the shortest-distance bin.

We again considered the same three hypotheses for cohesin–CTCF backside interactions (passing, blocking, and releasing), but with the CTCF position fixed and the enhancer position varied (20 evenly spaced values between 5 and 160 kb). We used p_c,f_ = 0.6 and p_c,b_ = 0 for the passing model, p_c,f_ = p_c,b_= 0.6 for the blocking model, and p_c,f_ = 0.6 and p_c,b_ = 0 together with p_u,b_ = 0.6 for the releasing model.

#### 3D polymer simulation

Three-dimensional chromatin conformations were simulated using the polychrom framework (OpenMM CUDA backend), following the loop-extrusion–polymer coupling protocol described in previous work. Simulations were performed using a Langevin integrator with error tolerance error_tol = 0.003, collision rate 0.03, single-precision arithmetic, and periodic boundary conditions in a cubic box at volume fraction 0.4.

The polymer backbone was modeled using harmonic bonds (bondLength = 1, bondWiggleDistance = 0.1), an angular potential (k = 1.5), and short-range excluded-volume interactions (polynomial_repulsive, trunc = 1.5, radiusMult = 1.05). Cohesin-mediated loops were represented as dynamic harmonic bonds between the two legs of each extruder (smcBondDist = 0.5, smcBondWiggleDist = 0.2). These bonds were updated at each cohesin step using a bond-updating scheme in which bonds corresponding to active loops were assigned finite stiffness, while inactive bonds were set to zero stiffness.

Loop extrusion dynamics were interleaved with polymer dynamics: each cohesin update step was followed by 5000 molecular dynamics steps. Polymer conformations were saved every 10 cohesin steps. To avoid excessive accumulation of inactive bonds, the simulation was reinitialized every 100 cohesin steps using the current conformation. The first 50 saved blocks were discarded as burn-in prior to analysis.

#### System replication for computational efficiency

To optimize GPU performance, simulations were performed on systems of approximately 100,000 monomers. This was achieved by tiling the loop-extrusion system into M independent copies of a smaller unit of size N:

● Single-CTCF simulations: N = 1_000, M = 100. Each unit contains a single CTCF site at position N/2, with independent loop-extrusion dynamics. The polymer is represented as a single continuous chain of N \times M = 100_000 monomers, such that adjacent units are connected along the backbone but evolve independently with respect to loop extrusion.
● Enhancer–promoter (EP) simulations: N = 500, M = 200. Each unit contains a full enhancer–promoter configuration, with the promoter at N/2 and the enhancer at N/2 + 160.

Contact maps were computed over individual units (length N for single-CTCF and 2N for enhancer-promoter configurations), and then averaged across all replicated units to obtain the final contact maps used for analysis and visualization.

#### Mapping between genomic and physical units

Each monomer corresponds to 1 kb of genomic distance and approximately 22 nm in physical space, based on calibration to experimental mean-squared displacement measurements. Contact maps were computed using multiple capture radii (specified in monomer units), corresponding to physical distances of 22 x cutoff nm.

#### Parameter sweep

For each condition, simulations were performed across a grid of extrusion parameters:

● Cohesin separation: d = 50, 100, 200 monomers
● Cohesin processivity: λ = 50, 100, 200, 400 monomers
● Contact radius: R_c_ = 2, 3, 5, 10 monomers (corresponding to 44, 66, 110, and 220 nm)

#### Quantification of contact enrichment

For each combination of (d, λ, R_c_) and condition, contact maps were computed by summing contributions across all replicated copies of the system (200 copies, each contributing a 2Nx2N window).

Promoter–enhancer contact frequency was extracted as the contact probability at genomic separation 160 monomers, corresponding to the matrix element: cmap[N/2; N/2 + 160].

To account for the baseline decay of contact probability with genomic distance, this value was normalized by the median contact frequency along the same diagonal (offset 160) within the same map. This yielded a distance-normalized contact enrichment.

To enable comparison across different parameter sets, enrichments were further normalized within each (d, λ, R_c_) combination by the corresponding CE2-only value, such that CE2-only = 1 by construction.

## Acknowledgements

We thank the Mirny and Nora labs, L. Giorgetti, E. de Wit, and their groups for thoughtful discussions. We are grateful to E. Anderson, F. Corsi, L. Giorgetti, M. Barbi, A. Hansen, A. Coulon, V. Ramani for comments on the manuscript. We thank L. Giorgetti for openly sharing unpublished data and analyses. We are very grateful to members of the UMR3664 at the Institut Curie for discussions and for hosting E. Nora during visits. AI disclosure: chatGPT4o and Claude were used as a grammatical assistant and coding aid; all resulting text was reviewed and further edited by humans.

## Author contributions

T.F. jointly with M.I. and H.D.P., developed the notion of cohesin bridging, developed analytical and computational models, performed computer simulations, comparison to experiments, parameter sweeps, developed figures and wrote the manuscript. M.I. performed 3D simulations and analysis of Hi-C data. LM. conceived the model of cohesin bridged encounters. L.M., jointly with M.I, G.F, N.A. conceived and supervised development of the model. L.M. N.A., G.F, M.I. studied the paradox of insufficient insulation in Hi-C, and developed the notion of cohesin-bridging as a path to its solution; realized the effectiveness of CTCF in facilitation. Genome engineering experiments were designed by K.H. and E.P.N. with input from T.F. and L.M., and K.H. performed all experiments with support from I.C. and T.H.; F.V.R. shared essential knowledge. T.F. and L.M. wrote the initial draft of the manuscript and together with K.H. and E.P.N. compiled data, assembled figures and edited the manuscript with input from all authors.

## Supplementary Figures

**Figure S1.**
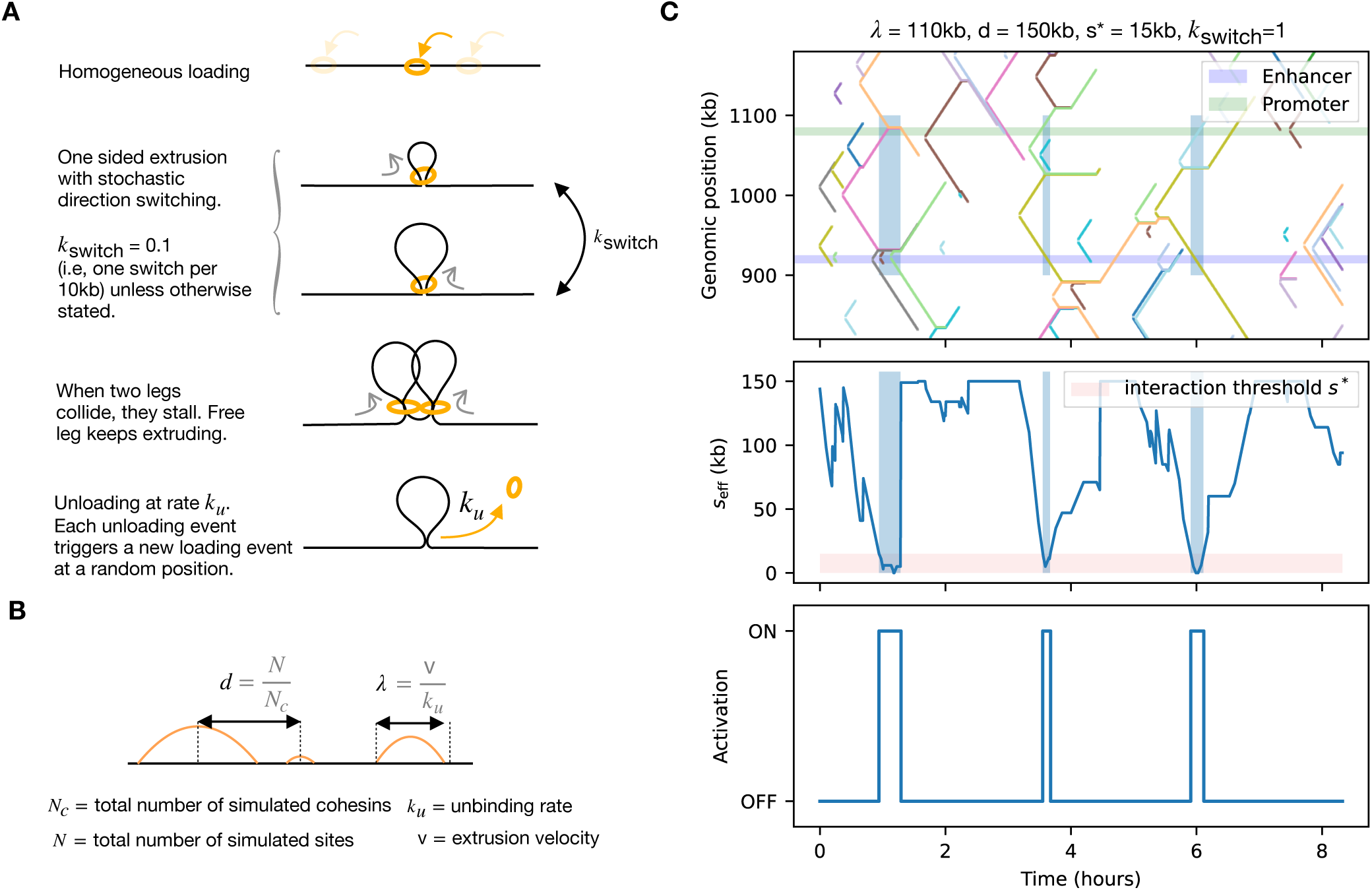
Simulation framework for cohesin-bridged model. (A) Schematic depiction of cohesin dynamics implemented in simulations. (B) Two key parameters characterize the steady state of the system. The average separation between loaded cohesins 𝑑, equal to the inverse linear density of cohesins. Second, the processivity *λ* is the average loop length of an unobstructed cohesin. (C) Example simulation of cohesin-bridged model. Top: kymograph showing the positions of loop-extruding cohesins over time, with the enhancer (green) and promoter (blue) marked as shaded regions, vertical blue shaded regions correspond to bridging events; (middle) the corresponding shortest genomic path 𝑠_+22_ connecting enhancer and promoter, shaded region corresponds to the linear distance below which the enhancer activates the promoter, *s\**; (bottom) the promoter activation state, which switches to *ON* whenever

**Figure S2.**
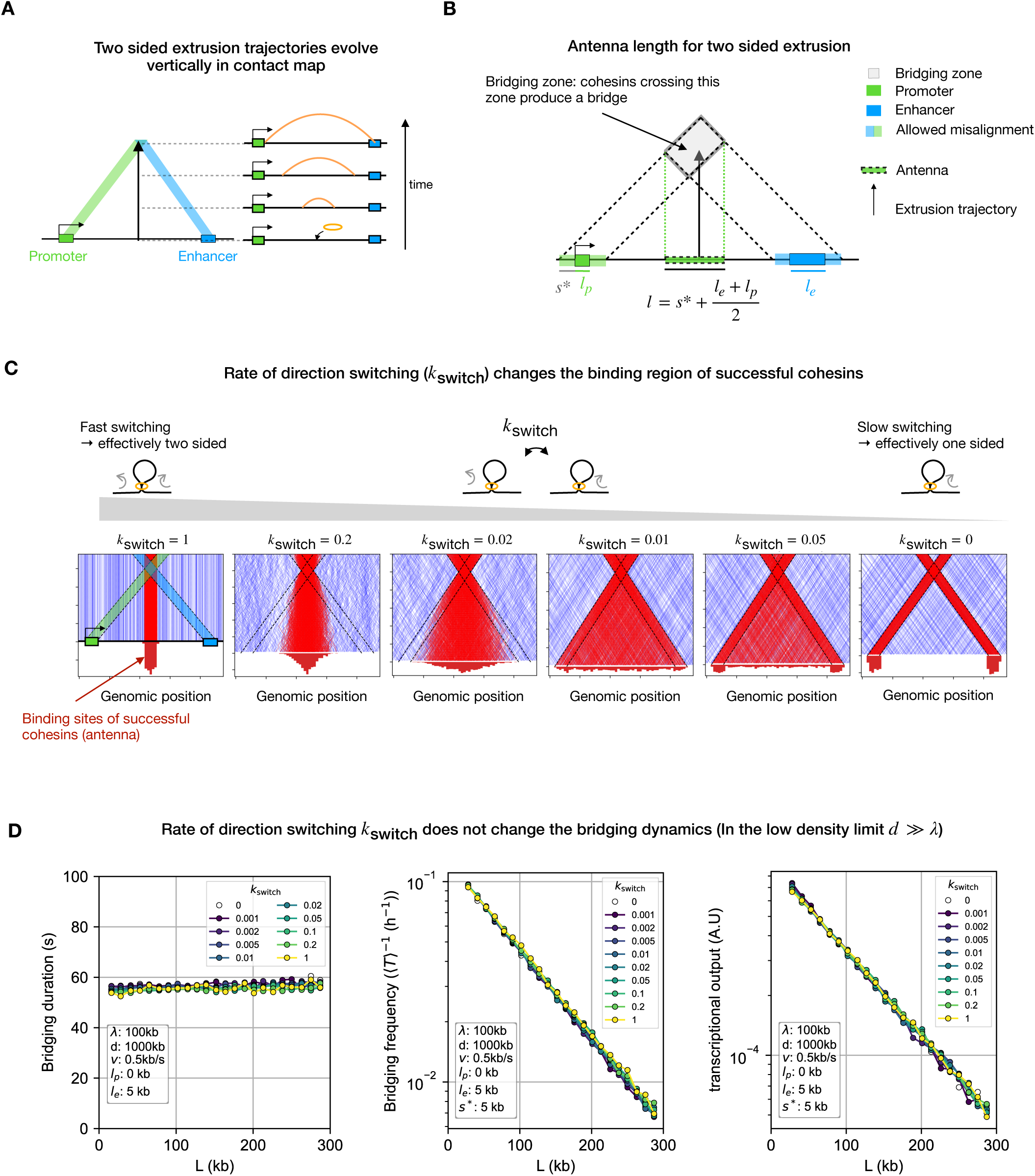
The rate of extrusion direction switching modifies antenna position and length but not bridging dynamics. (A) Explanatory schematic for extrusion trajectory in a contact map. Each line represents the trajectory of a single cohesin in a contact map. At every moment in time, a cohesin holds two chromatin sites (one point in the contact map); connecting all pairs of chromatin sites that a cohesin has held during its residence time forms a continuous track within the contact map. Each track starts at the bottom because cohesin initially connects nearby sites and moves upwards as the loop enlarges and connects more distant sites. The colored lines emanating from the enhancer and promoter represent their bridging zones, i.e., regions of the map that connect to the enhancer (blue) and promoter (green). Cohesins that pass by their intersection (arrowhead) have successfully bridged the enhancer and promoter. (B) Geometry underlying the calculation of antenna length for two-sided extrusion. The antenna corresponds to the binding sites that allow for enhancer-promoter bridging, i.e., from where on the x-axis (where cohesin trajectories start) cohesins can access the bridging zone (grey region). For two-sided extruders, which are vertical lines in the contact map representation, this amounts to projecting the bridging zone onto the x-axis vertically. Simple geometry then shows that this corresponds to a central region of length 𝑠 ∗ +(𝑙. + 𝑙_+_)/2. See supplementary text for more details. (C) Simulated extrusion trajectories illustrating the dependence of the antenna on switching rates. Contact maps show extrusion trajectories for one-sided extruders at different direction-switching frequencies, with spatial distribution of loading positions of cohesins that successfully bridge the enhancer and promoter (red histogram on the bottom). Left: at high switching frequency, extrusion is effectively two-sided, and successful trajectories (red) originate from the central region. Right: at zero switching frequency, extrusion is strictly one-sided, and successful trajectories originate from regions near the enhancer and promoter. As the switching frequency varies between these extremes, the antenna continuously deforms and shifts between these two configurations. Note that, for these visual representations, a simplified simulation was used where cohesins are not blocked by collisions with other cohesins. (D) Bridging dynamics are insensitive to direction switching frequency in the low cohesin density limit. Bridge frequency (Left), duration (middle) and transcriptional output (total time in bridged state, right) as a function of enhancer-promoter distance in simulations for different direction switching frequencies (kswitch).

**Figure S3.**
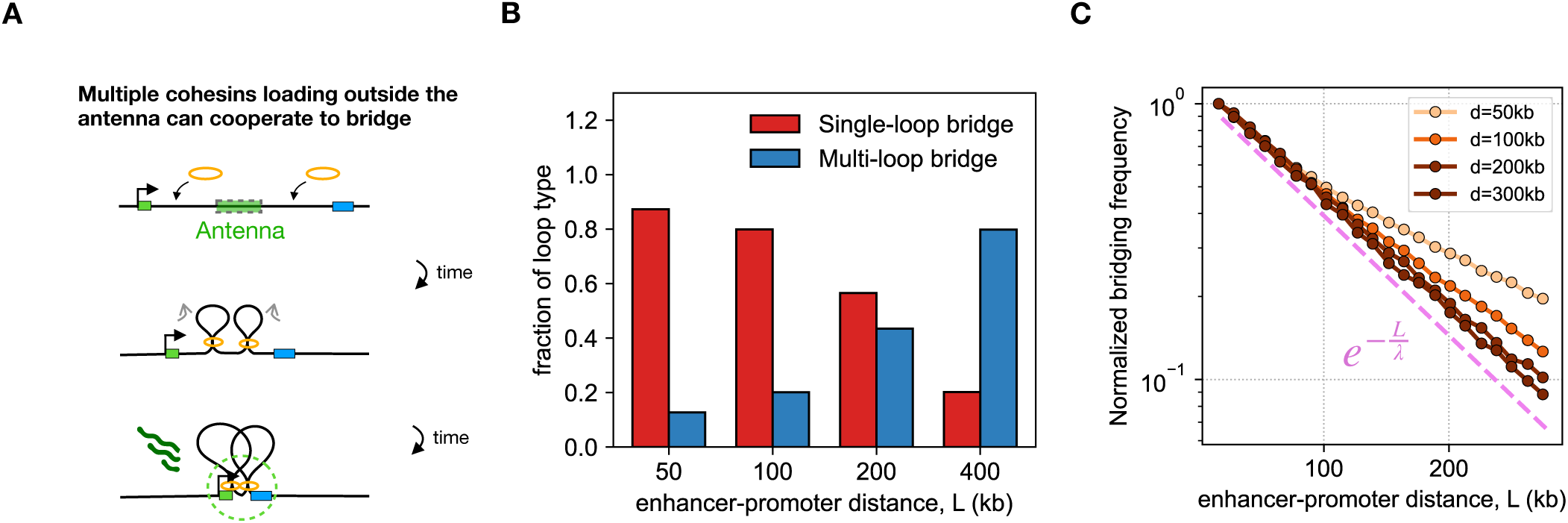
Multi-loop bridging by cohesin extends long-range enhancer–promoter interactions. **(A)** Schematic showing how two cohesins that land outside the antenna can cooperate to bridge the enhancer and promoter. **(B)** Fraction of single loop-bridging (red, bridges formed by one cohesin) vs multi-loop bridging (blue, bridges formed by more than one cohesin) as a function of enhancer-promoter separation, *L* (*d* = 150 kb, *λ* = 110 kb). The fraction of multi-loop bridges becomes dominant when *L* >> *d*. **(C)** Simulated enhancer–promoter bridging rate as a function of genomic distance *L,* for different cohesin separations (*d*). For non-interacting cohesins (grey, analytical theory), the decay follows a simple exponential ∼ 𝑒 ^-L/λ^. Simulating interacting cohesins (where collided legs stop extruding–brown curves, simulations) yield shallower decays of enhancer-promoter bridging rates, reflecting cooperative bridging through multi-loop conformations that provide added bridging opportunities–especially at larger distances.

**Figure S4.**
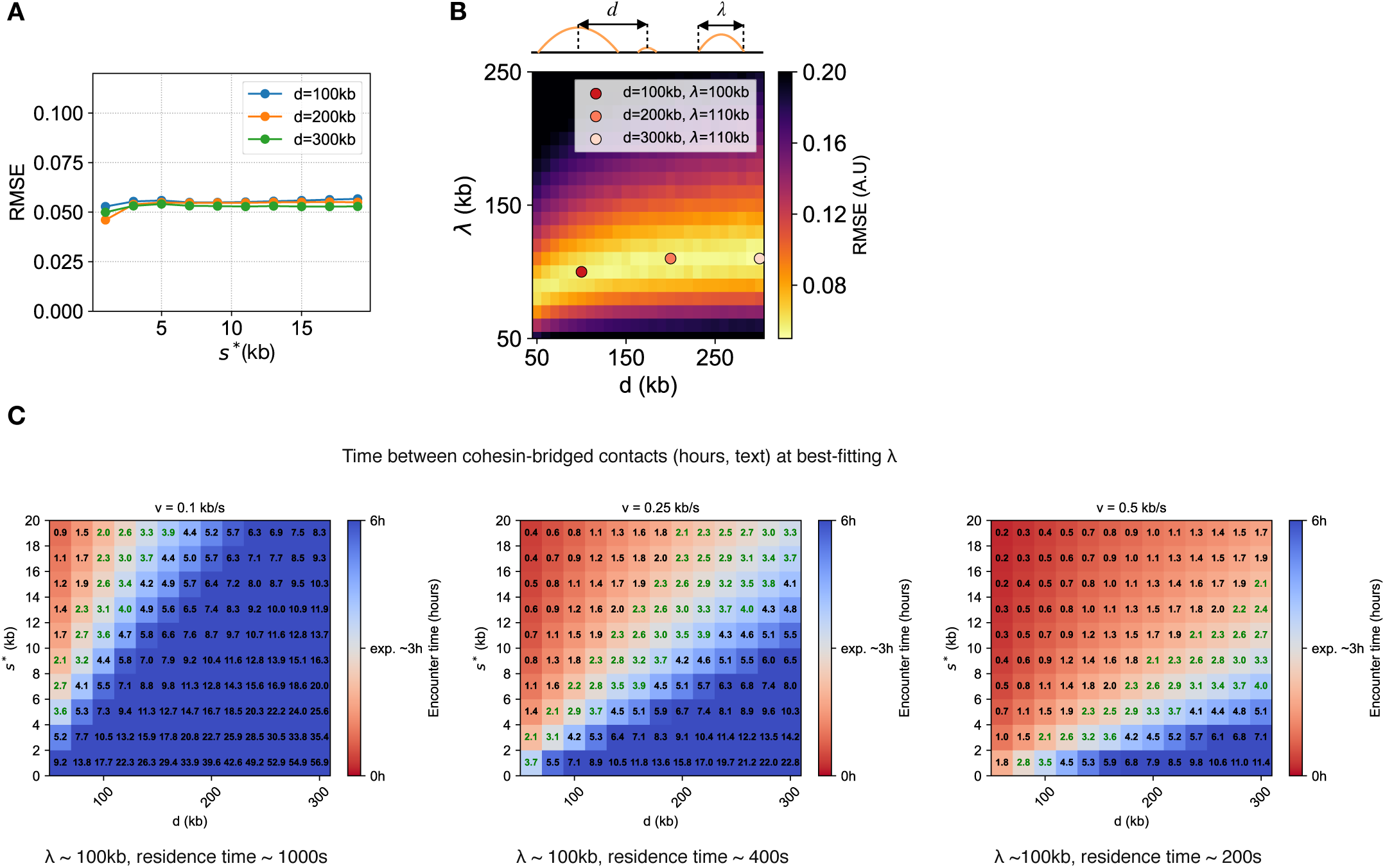
Fitting cohesin-bridging model to Sox2–SCR distance dependence. **(A)** Best-fit Root Mean Square Error (RMSE) as a function of interaction threshold *s\** for different cohesin separations *d* (optimized over cohesin processivity *λ*). The weak dependence of RMSE over *s\** indicates that the quality of the fit to experimental data is largely insensitive to *s\**. This reflects the fact that *s\** only changes the prefactor of the decay but not the rate of decay, which is controlled by *λ*. **(B)** RMSE (color: yellow corresponds to better fit) between the predicted transcription levels and the Sox2 integration data, for a grid of cohesin processivities *λ* and separations *d* at *s\**=10kb. Dots correspond to best fitting (*λ, d*) pairs for four different values of *d* (100, 150, 200, 300kb). **(C)** For each extrusion velocity, heatmaps show the predicted time between bridges (text and color) as a function of cohesin separation d and interaction threshold *s\** at best fitting value of λ. The colormap is centered on the experimental estimate of 3 hours (*52*), making the best fitting values appear in white. Numbers in green indicate predictions within ± 1 h of this value.

**Figure S5.**
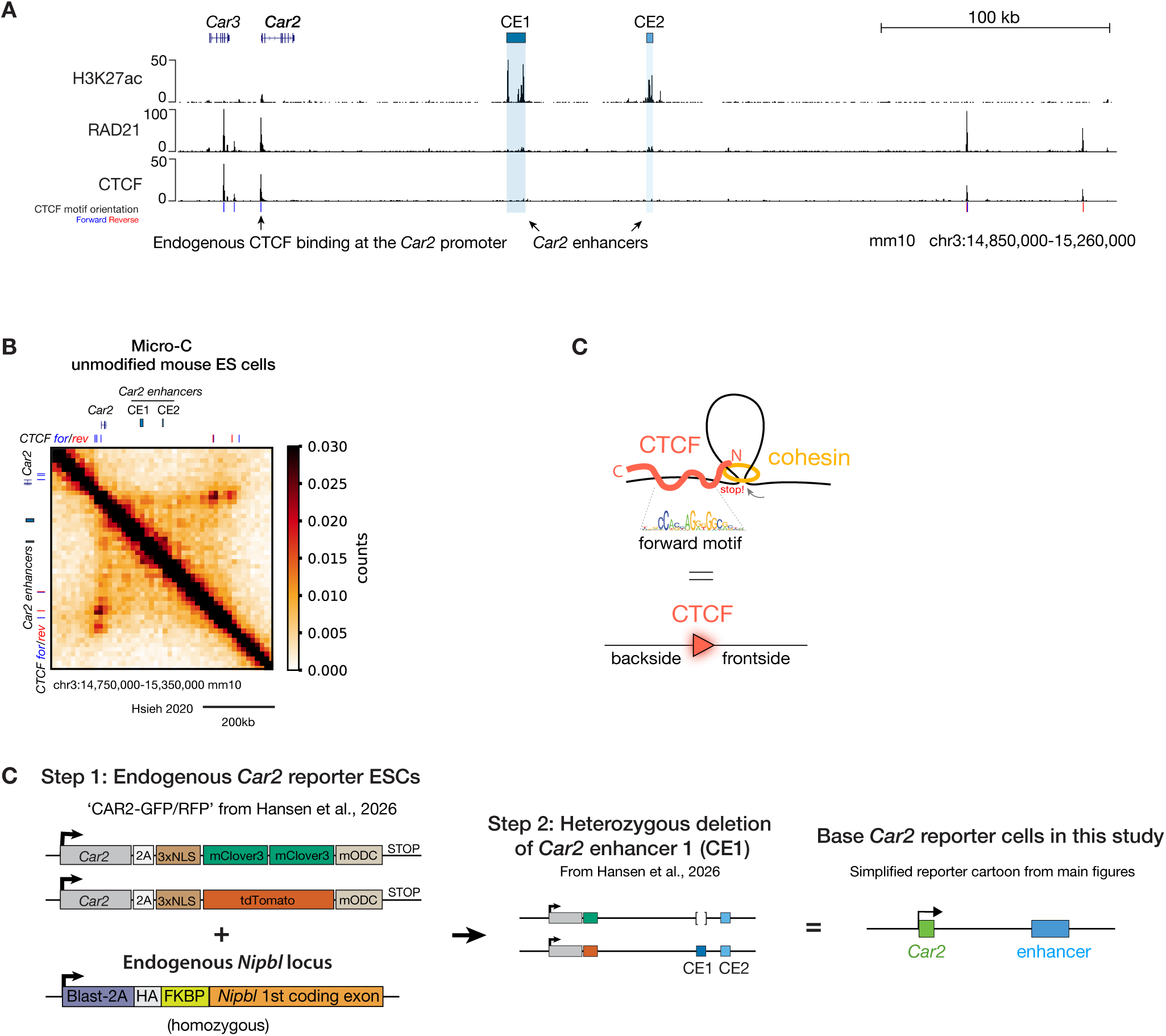
Details of the *Car2* locus and reporter line used for all experiments. (A) Genome browser view of the *Car2* locus displaying related CUT&Tag for H3K27ac (*33*), and ChIP-seq for RAD21 and CTCF (*93*) in ESCs. Forward (blue) and reverse (red) CTCF motif orientation is indicated. *Car2* enhancer 1 (CE1) and *Car2* enhancer 2 (CE2) are highlighted in blue. (B) Micro-C contact map of the *Car2* locus in unmodified mouse ESCs (*74*). (C) Cartoon explaining the orientation nomenclature of CTCF sites. When a forward CTCF motif faces incoming cohesin, the N-terminus of CTCF engages it, blocking and retaining cohesin, thereby stabilizing the DNA loop. Frontside and backside encounters correspond to cohesin approaching the N-terminus (N) or C-terminus (C) of the CTCF protein, respectively. (D) Details of the *Car2* fluorescent reporter ESCs. The endogenous *Car2* locus was previously edited to express a 2A-2XmClover3 (GFP) cassette from one allele and a 2A-tdTomato (RFP) cassette from the other allele, in NIPBL-FKBP degron ESCs (*33*). To simplify interpretations, a cell line with one enhancer (CE1) removed was used, leaving only CE2. A simplified schematic of the resulting locus (right) is used in all main and supplementary figures.

**Figure S6.**
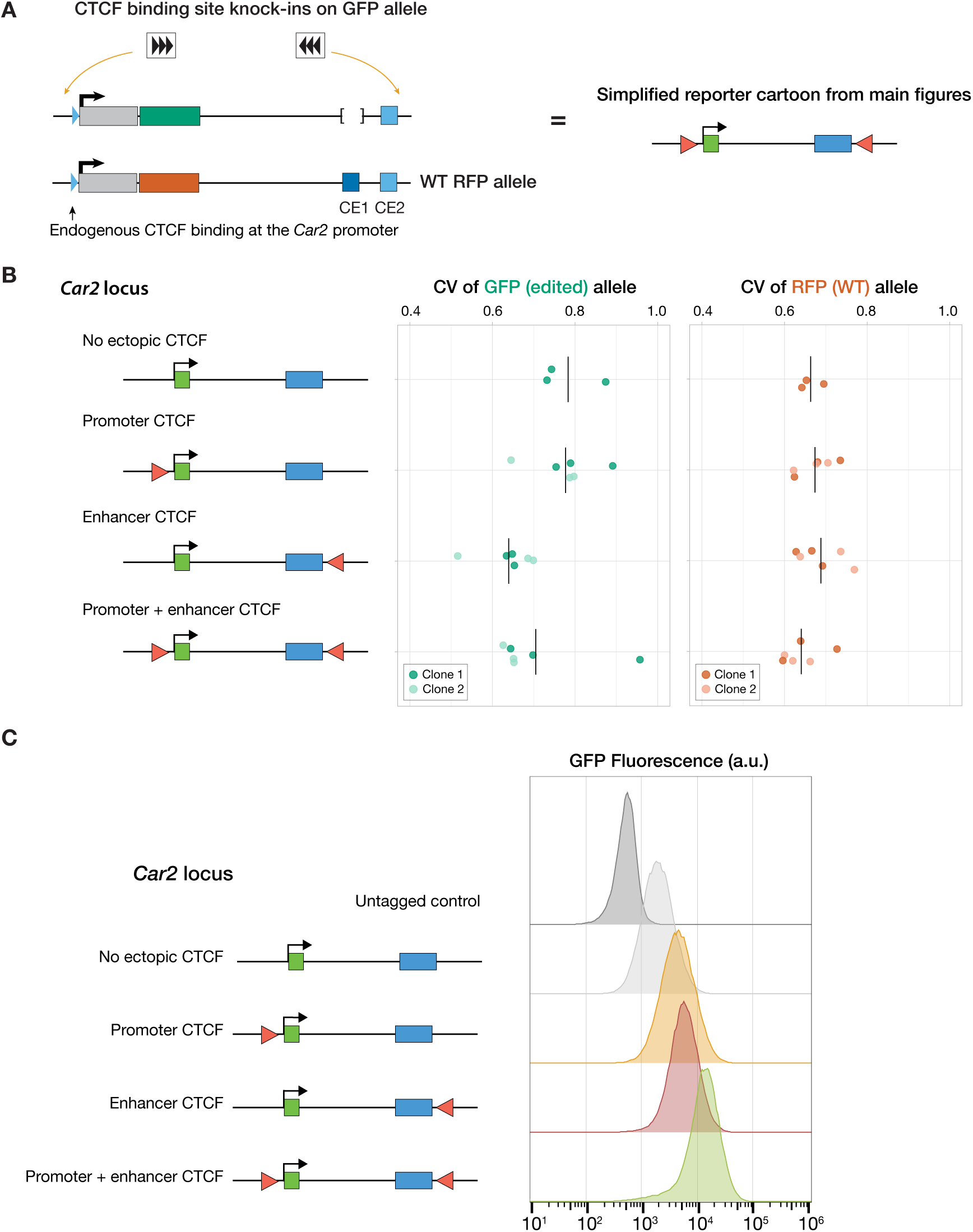
Supporting analysis of facilitation by CTCF at the *Car2* locus. **(A)** Details for the simplified schematic of the *Car2* reporter with insertions of 3X facilitating CTCF sites at the promoter or enhancer (CE2). **(B)** Coefficient of Variation (CV) for each layout, colored by individually derived clone. CV is calculated as the ratio of the standard deviation to the mean of the population. Notch indicates the average across all clones (n = 3 flows each for 2 clones). **(C)** Representative flow histograms of clone 1 from (B), showing a unimodal distribution of *Car2* reporter expression across single cells.

**Figure S7.**
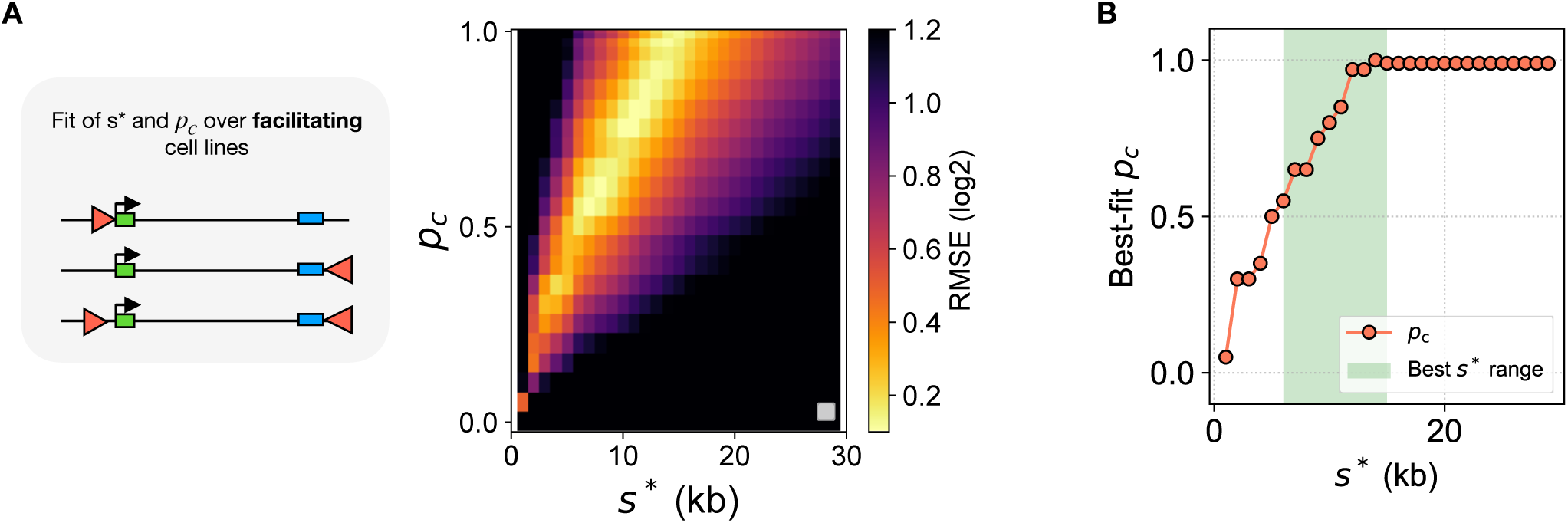
Parameter fitting for facilitation at the *Car2* locus. **(A)** log2 RMSE between CTCF layouts involving facilitating CTCF and simulations as a function of the threshold *s\** and the CTCF capture probability *pc*. **(B)** Best fitting *pc* as a function of *s\**. Green shaded region represents values of *s\** for which the RMSE at best *pc* is lower than 0.2 (4 – 16 kb).

**Figure S8.**
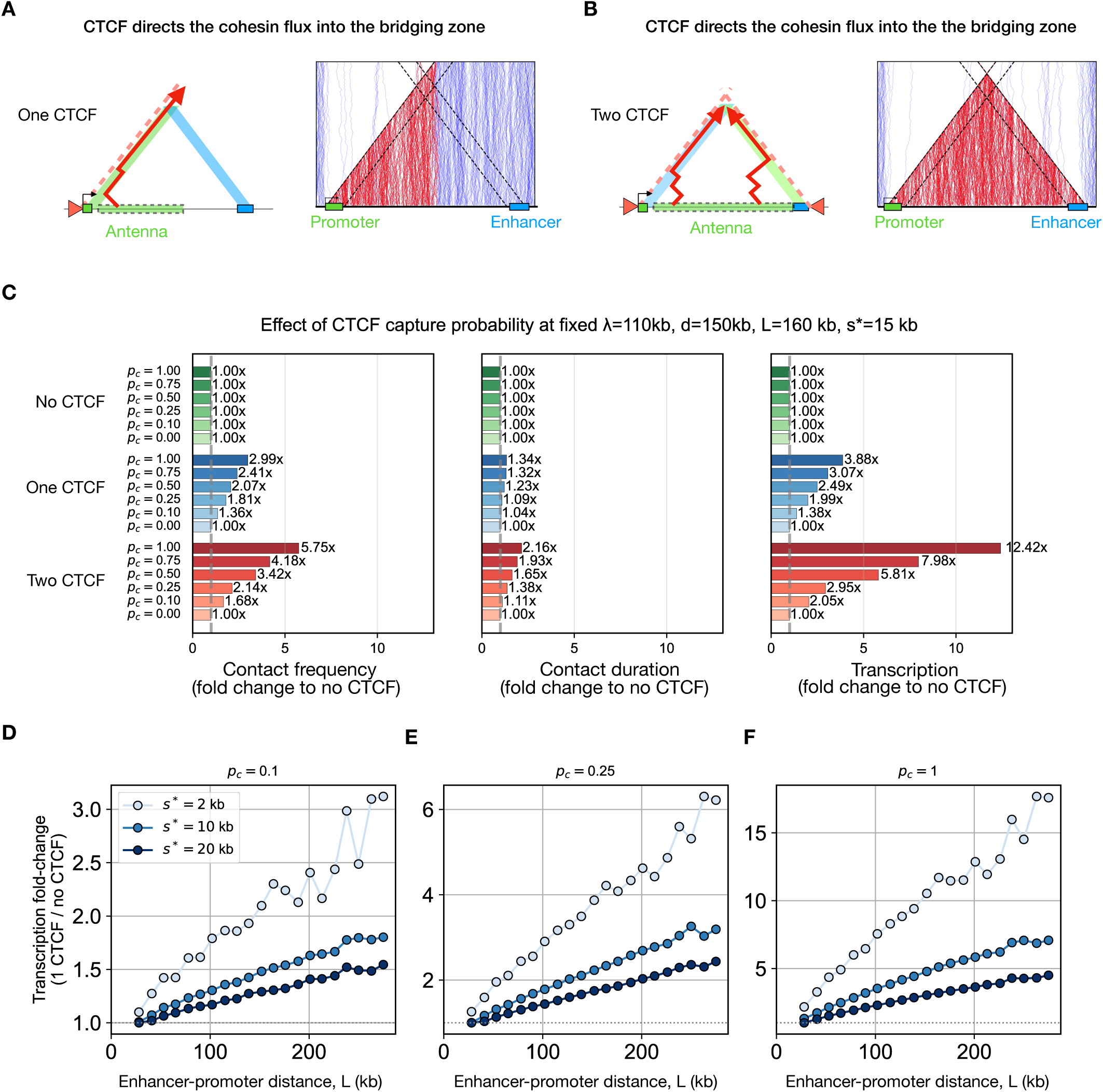
CTCF at the enhancer or promoter increases transcriptional output. (A) CTCF binding at the promoter or enhancer redirects the cohesin flux to the bridging zone. Left: schematic showing a cohesin trajectory (red arrow) that would fail to bridge in the absence of CTCF but is redirected into the bridging zone by CTCF. This extends the antenna (green box) to half of the enhancer-promoter region. Right: corresponding simulation (see fig. S2C caption for simulation details). (B) Same as (A) for CTCF at both the enhancer and promoter. (C) Effect of CTCF sites on bridging frequency, duration and transcriptional output function of *pc*. (D) Transcriptional fold change upon insertion of a single CTCF site as a function of enhancer–promoter distance. Fold change (relative to the no-CTCF condition) is shown for different values of the activation threshold (*s*\*), for CTCF capture probabilities (*p*c=0.1). For small *s\** at 200kb even *pc* = 0.1 leads to ∼2 fold increase in transcription. (E) Same as (D) for *p*c=0.25 (F) Same as (D) for *p*c=1

**Figure S9.**
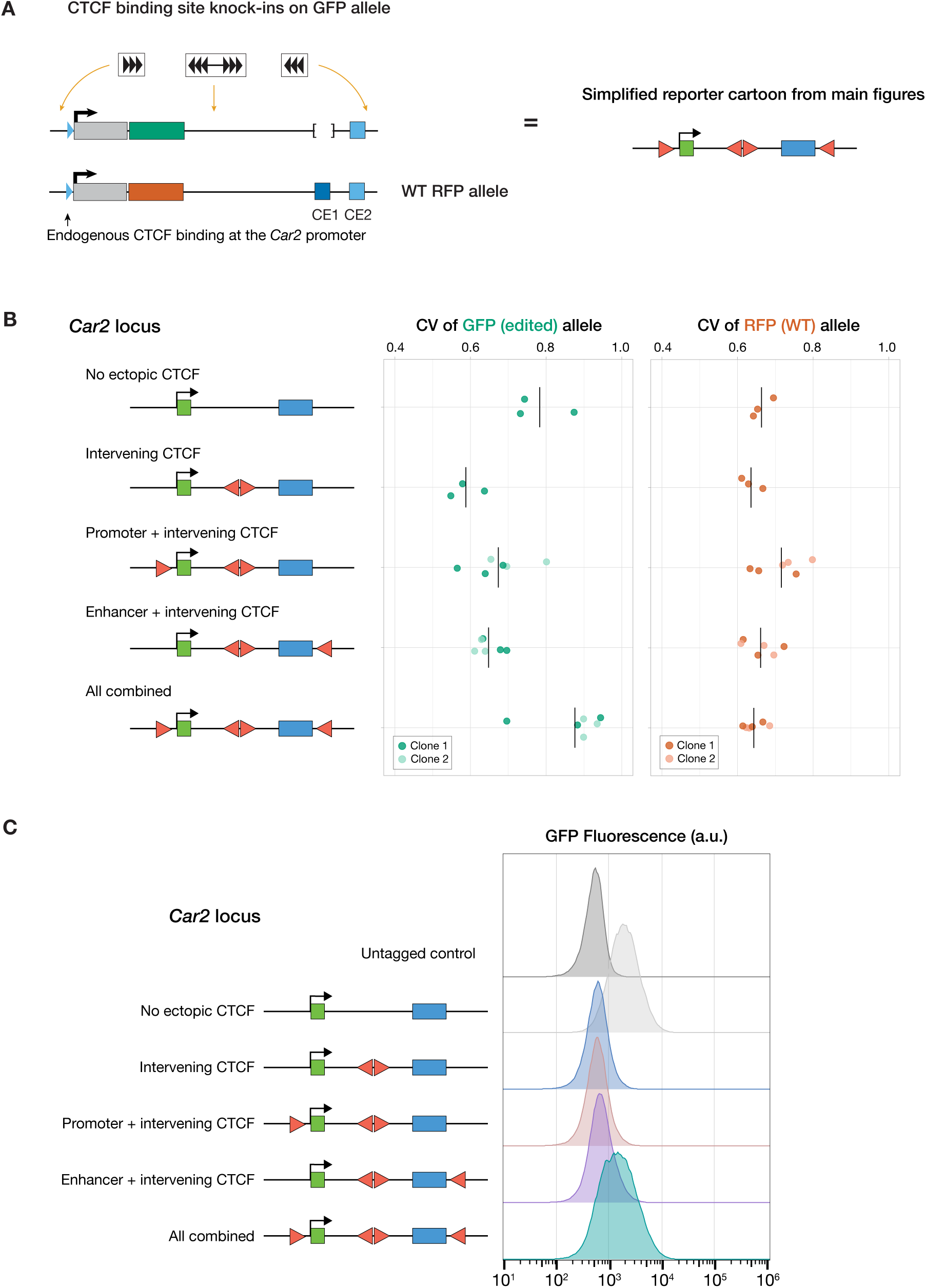
Supporting analysis of insulation by CTCF at the *Car2* locus. (A) Details for the simplified schematic of the *Car2* reporter with insertions of 6X divergently-oriented insulating CTCF sites between *Car2* and CE2, with or without facilitating CTCF sites. (B) Coefficient of Variation (CV) for each layout, colored by individually derived clone. Notch indicates the average across all clones (n = 3 flows each for 2 clones). Note, the second clone for Intervening CTCF was not included because expression levels were below detectable limits. (C) Representative flow histograms of clone 1 from (B), showing a unimodal distribution of *Car2* reporter expression across cells.

**Figure S10.**
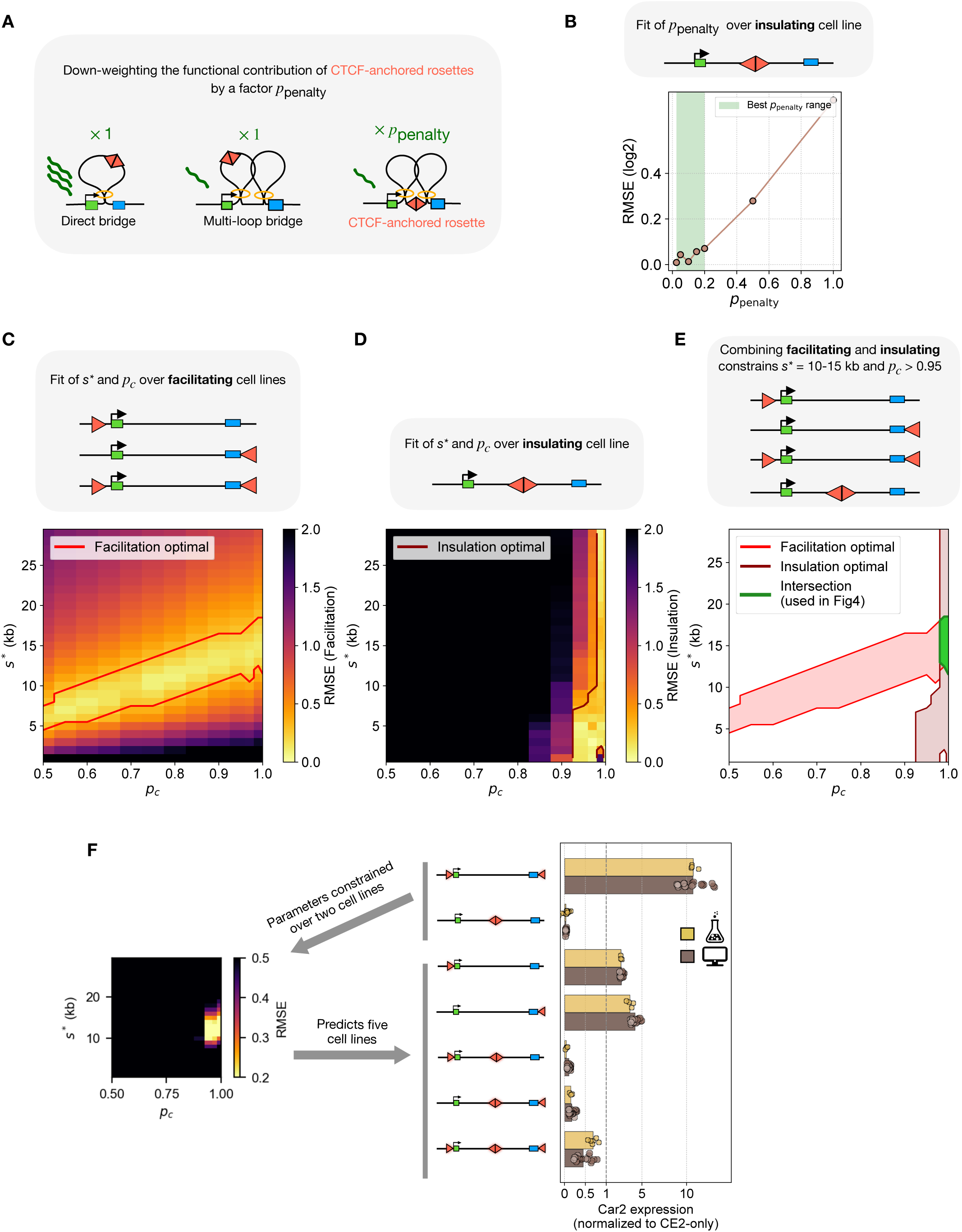
Fitting parameters to facilitation and insulation by CTCF at the *Car2* locus. (A) Schematic of contribution of different bridging types to transcription. All bridging types are considered to contribute equally to transcription (left and middle), except for bridging through CTCF-anchored rosettes (right) that are down-weighted by a factor *ppenalty* (see Methods). Here multi-loops resulting from simple cohesin collisions (middle) are considered to contribute normally to transcription. (B) log2 RMSE between insulation cell line (6x CTCF cassette at +58kb from the promoter) and corresponding simulation function of *ppenalty*. Green shaded region corresponds to values of *ppenalty* where RMSE < 0.1 (**C-E**) Fitting of both facilitating cell and insulating cell lines constrain *s\** and *pc*. (C) Heatmap of RMSE between all facilitating cell lines and corresponding simulation function of *pc* and *s\**. Contoured region correspond to parameter space where RMSE < 0.2. This fit constrains the parameters to a family of solutions but does not uniquely determine both parameters. (D) Heatmap of RMSE between insulating cell line and corresponding simulation function of *pc* and *s\**. Contoured region correspond to parameter space where RMSE < 0.4. The strong insulation observed in this configuration constrains *pc* to be larger than 0.95. (E) Intersecting both insulating and facilitating layouts constrains the parameters to a small region of parameter space. (F) Parameter inference from minimal data. Fixing *ppenalty* = 0.2 from the insulating cell line, other parameters (*s\** and *pc*) can be inferred using only two conditions: one facilitating configuration (e.g., CTCF at both enhancer and promoter) and the insulating configuration. These parameters are then used, without further fitting, to predict transcriptional output across all other configurations. Each point corresponds to a distinct simulation and bars correspond to average prediction over these simulations. Experimental datapoints reproduced from Fig. 4.

**Figure S11.**
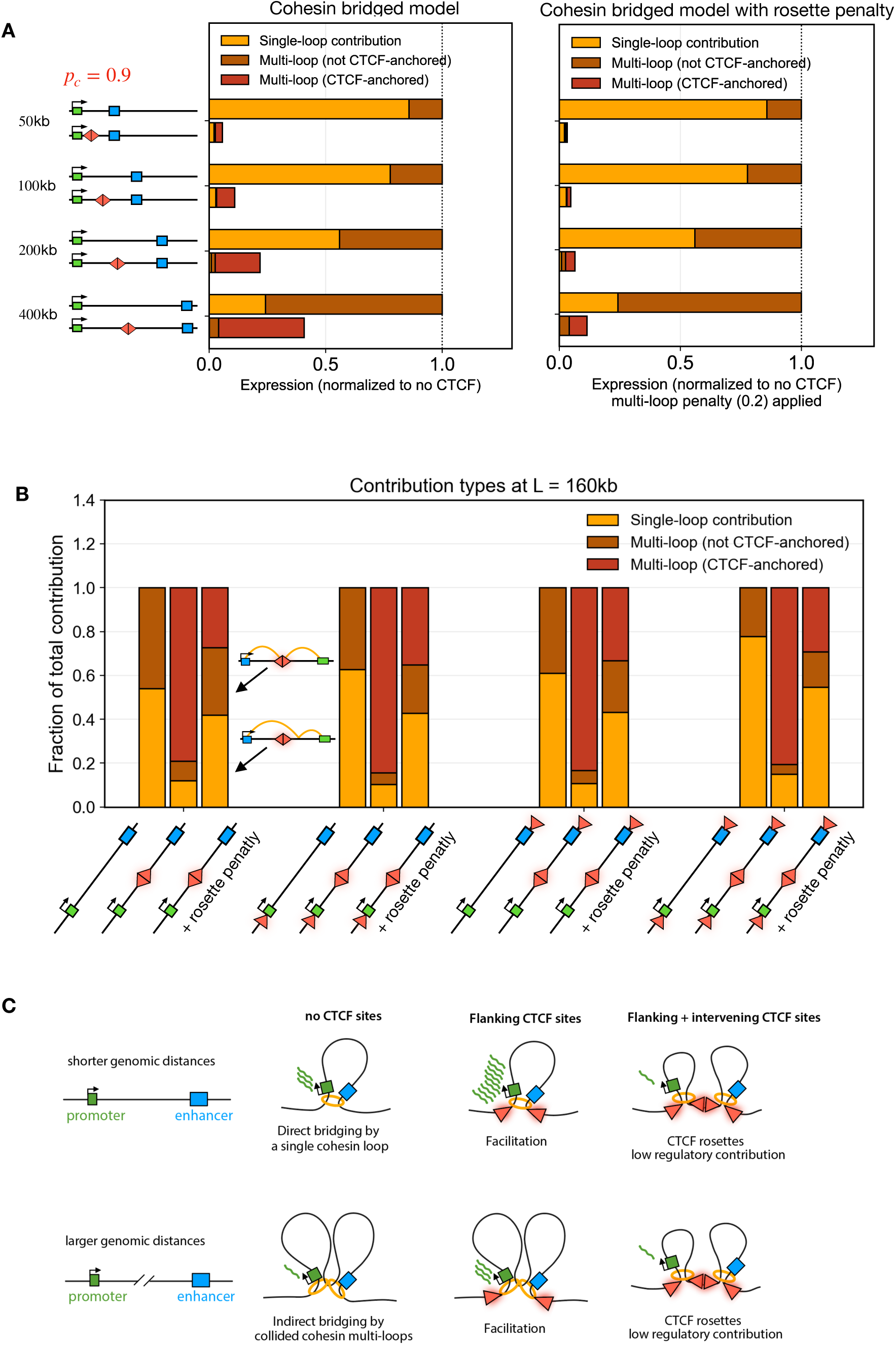
Functional contribution of different bridging types. (A) Contribution of bridging types (single-loop, multi-loop, CTCF-anchored rosettes) to transcriptional output, with and without intervening CTCF sites, for different enhancer-promoter separations. Simulations in the left panel consider all bridge types can contribute equally to transcription. Simulations in the right panel consider that CTCF-anchored rosettes contribute 0.2 times the other types of bridges. At short distances, in the absence of CTCF, most bridging events are single loops; intervening CTCF sites prevent these bridges and cause ∼10 fold transcriptional insulation–even without downweighting CTCF-anchored rosettes. As distance increases, the fraction of multi-loop increases; most of these multi-loop bridges remain in the presence of the intervening CTCF sites as CTCF-anchored rosettes. Therefore, to explain the strong transcriptional insulation by CTCF at large distances observed experimentally, CTCF-anchored rosettes must contribute less to enhancer-promoter communication than other types of bridges. (B) Same as (A) but at fixed enhancer-promoter distance and for different CTCF layouts, with and without the CTCF-anchored downweighting. Total transcription was normalized within each condition to show how the contribution of each type changes for different CTCF layouts and when the downweighting is applied. (C) Cartoon diagrams of the distinct types of enhancer-promoter bridges and their relative regulatory contribution across genomic distances.

**Figure S12.**
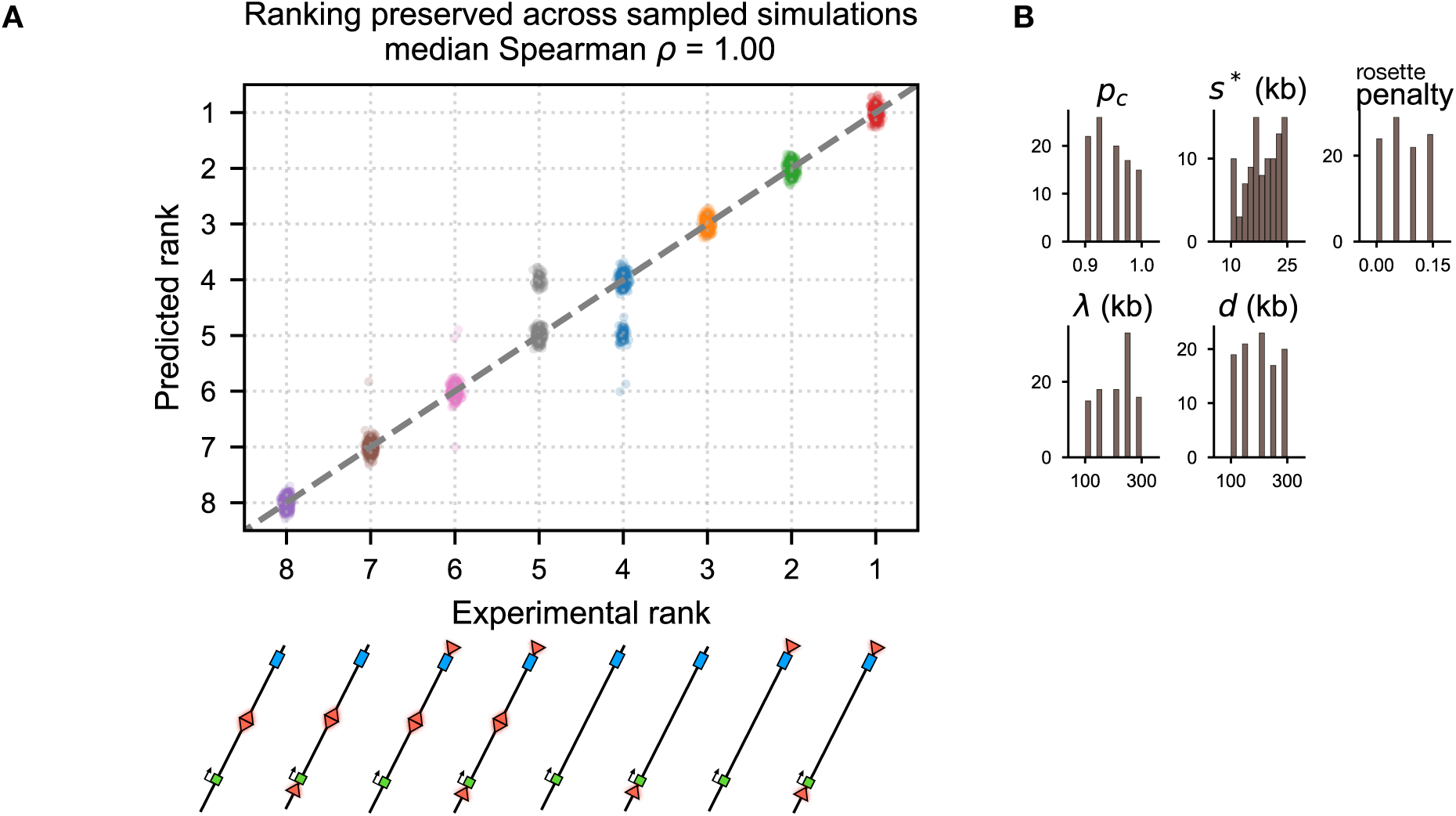
Model predictions robustly rank transcriptional outputs across CTCF layouts independent of precise parameter values. (A) Ranking agreement between experiments and model predictions across randomly sampled parameters. Each point corresponds to a single simulation parameter set, showing the rank of transcriptional output across cell lines in experiments (x-axis) versus simulations (y-axis). Ranks are computed from transcription values and ordered from highest to lowest. The dashed diagonal indicates perfect agreement between experimental and predicted rankings. Nearly every parameter set tested preserved the relative ordering of experimental transcriptional outputs across all CTCF layouts (median Spearman ρ = 1.00), indicating that insulation and facilitation robustly emerge from the mechanism of cohesin-bridging rather than from parameter tuning of the model. (B) Distribution of sampled parameter values used in (A).

**Figure S13.**
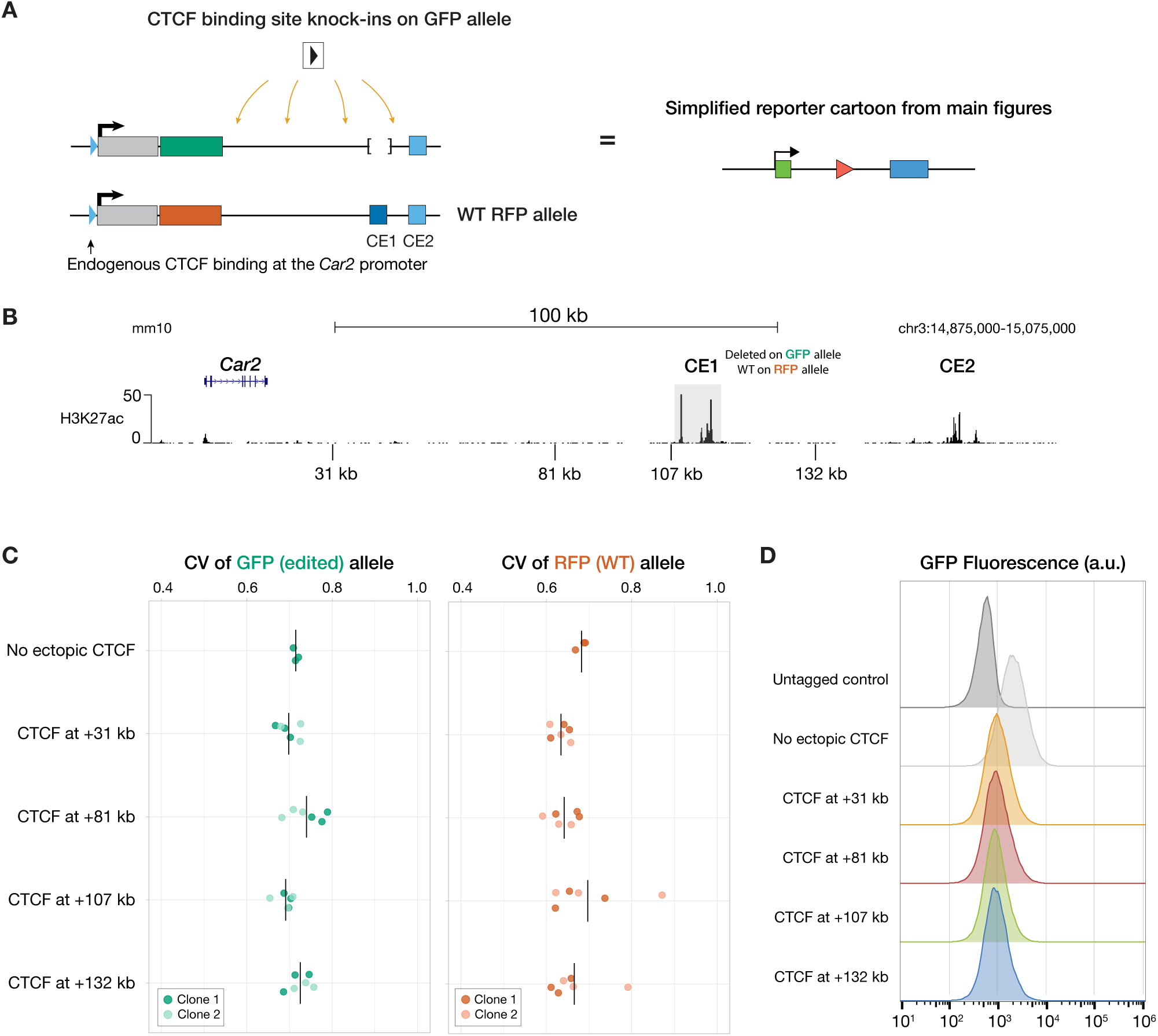
Supporting analysis of insulation by a single CTCF across the *Car2* locus. (A) Details for the simplified schematic of the *Car2* reporter with an insertion of a single CTCF site in the forward orientation at various positions between *Car2* and CE2. (B) Genome browser view of the *Car2* locus displaying related CUT&Tag for H3K27ac in ESCs (*33*) and annotated positions for CTCF site insertions. (C) Coefficient of Variation (CV) for each layout, colored by individually derived clone. Notch indicates the average across all clones (n = 3 flows each for 2 clones). (D) Representative flow histograms of clone 1 from (C), showing a unimodal distribution of *Car2* reporter expression across cells.

**Figure S14.**
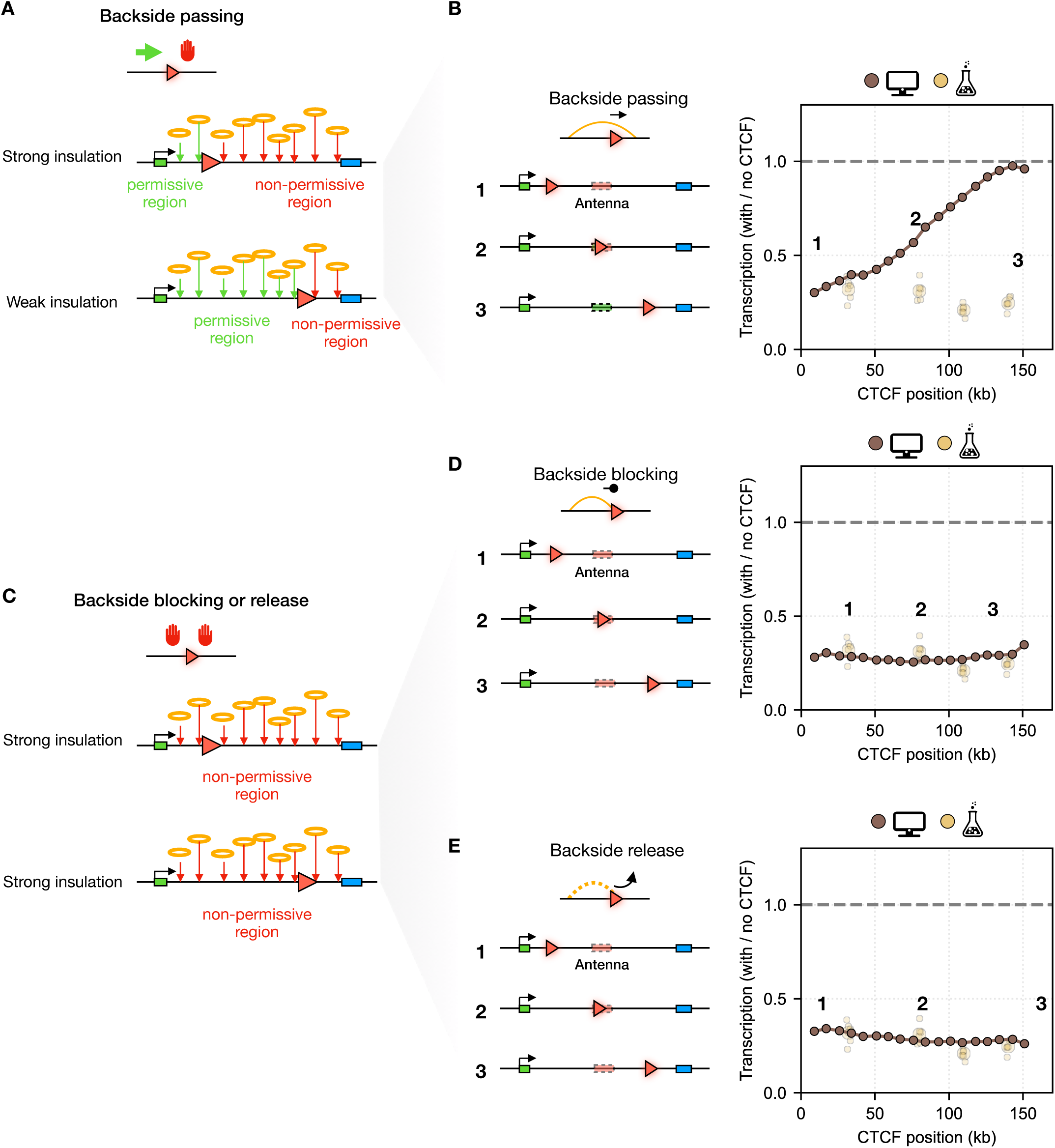
Three models of cohesin-CTCF backside encounters predict distinct insulation profiles. (**A**) Schematic illustrating how backside passing of CTCF sites partitions the enhancer–promoter region into a permissive region (green) and a non-permissive region (red). Cohesins approaching from the permissive side can traverse the site, whereas those approaching from the non-permissive side are blocked. (**B**) Left: Schematic depicting why insulation by a single CTCF site under the backside passing hypothesis produces position-*dependent* insulation. Right: predicted transcriptional output as a function of the location of the inserted CTCF site, normalized to the layout without ectopic CTCF site (same as Fig. 6E). (C) Schematic illustrating why insulation under the backside-blocking hypothesis produces position-*independent* insulation. Cohesin is blocked with equal probability from both sides, eliminating the permissive/non-permissive asymmetry. (D) Same as in (B), but under the backside-blocking hypothesis. This produces position-*independent* transcriptional insulation. (E) Same as in (B), but under the backside-release hypothesis, where backside encounters cause cohesin unloading or loop release rather than capture. This produces position-*independent* transcriptional insulation.

**Figure S15.**
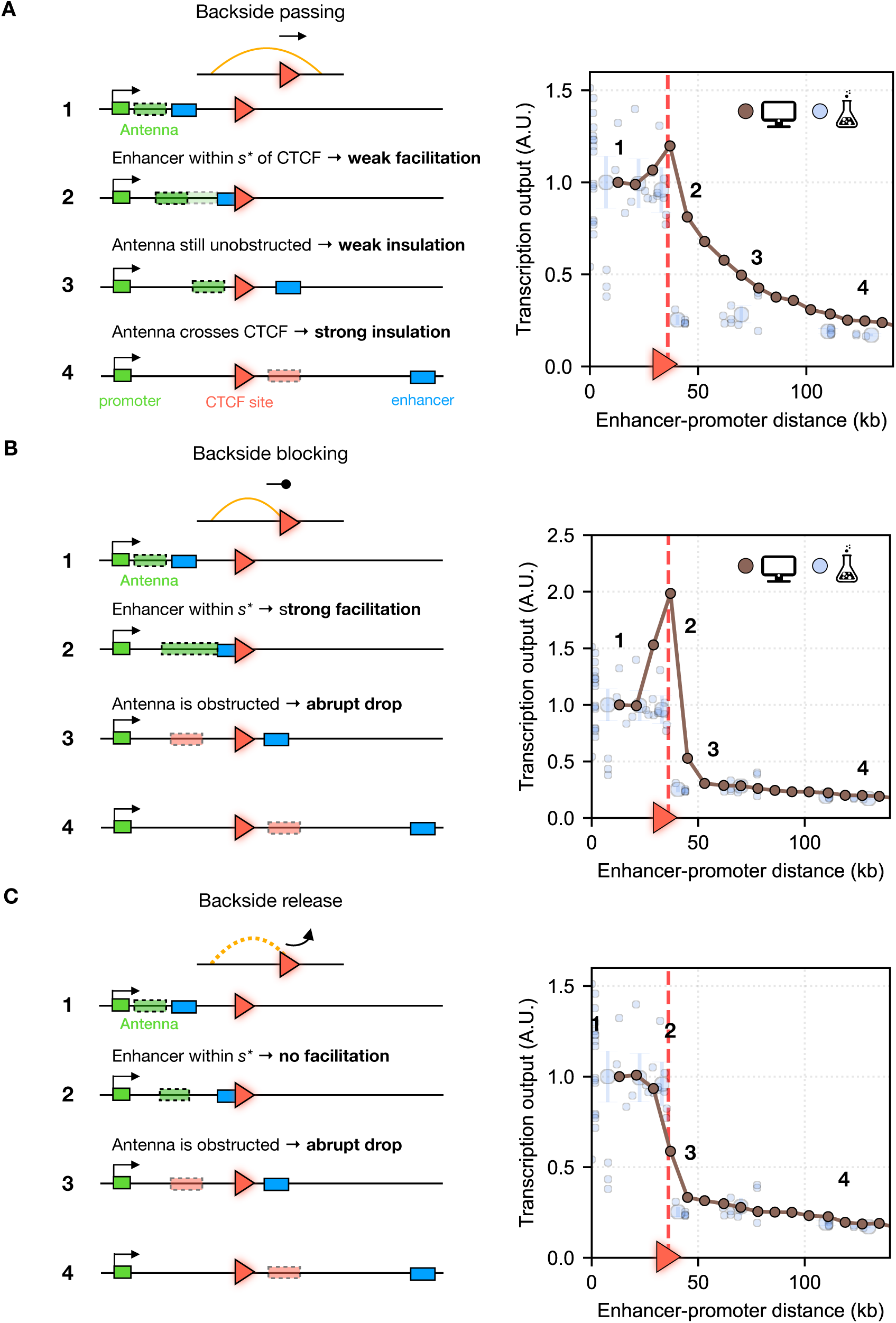
CTCF interaction rules determine distance-dependent transcriptional responses across a CTCF site at the *Sox2* reporter locus. For each panel, the enhancer is moved to increasing distances from the promoter across a fixed, forward-oriented CTCF site. Four regimes are illustrated as schematics (left) and as predicted versus observed transcriptional output (right; reproduced from Fig. 6; experimental data from (*26*)). **(A) *Backside-passing hypothesis***: cohesin passes freely through the backside of the CTCF site. (1) The CTCF site lies outside the enhancer–promoter interval: no effect on transcription. (2) The enhancer is within *s\** of the CTCF site: weak facilitation, because cohesins approaching from the backside occasionally stall when the site is already occupied by a frontside-blocked cohesin. (3) The enhancer is across the CTCF site but at short distance, such that the antenna remains on the backside: weak insulation. (4) Larger distances: the antenna is across the CTCF site; strong insulation, as productive cohesins loaded at the antenna now encounter the blocking side of the CTCF site. The backside-passing hypothesis therefore predicts a gradual decline in transcription as distance increases from CTCF site, inconsistent with the sharp drop observed experimentally. **(B) *Backside-blocking hypothesis***: cohesin is blocked by the CTCF site from both back-and frontsides. (1) The CTCF site lies outside the enhancer-promoter interval: no effect on transcription. (2) The enhancer is within *s\** of the CTCF site: strong facilitation due to backside blocking. (3) The enhancer is across the CTCF site: abrupt drop in transcription, cohesins coming from the antenna are blocked upon encountering the CTCF backside, before forming an enhancer-promoter bridge. (4) Larger distances: transcription remains strongly repressed. The backside-blocking hypothesis predicts spurious facilitation near the CTCF site (2), inconsistent with experiments. **(C) *Backside-release hypothesis***: backside encounters cause cohesin to release the DNA loop. (1) The CTCF site lies outside the enhancer-promoter interval: no effect on transcription. (2) The enhancer is within *s\** of the CTCF site: no facilitation, as backside release prevents loop retention. (3) The enhancer is across the CTCF site: abrupt drop in transcription, as cohesins loaded at the antenna become released before forming an enhancer-promoter bridge. (4) Larger distances: transcription remains strongly repressed. The backside-release model correctly reproduces experimental observations.

**Figure S16.**
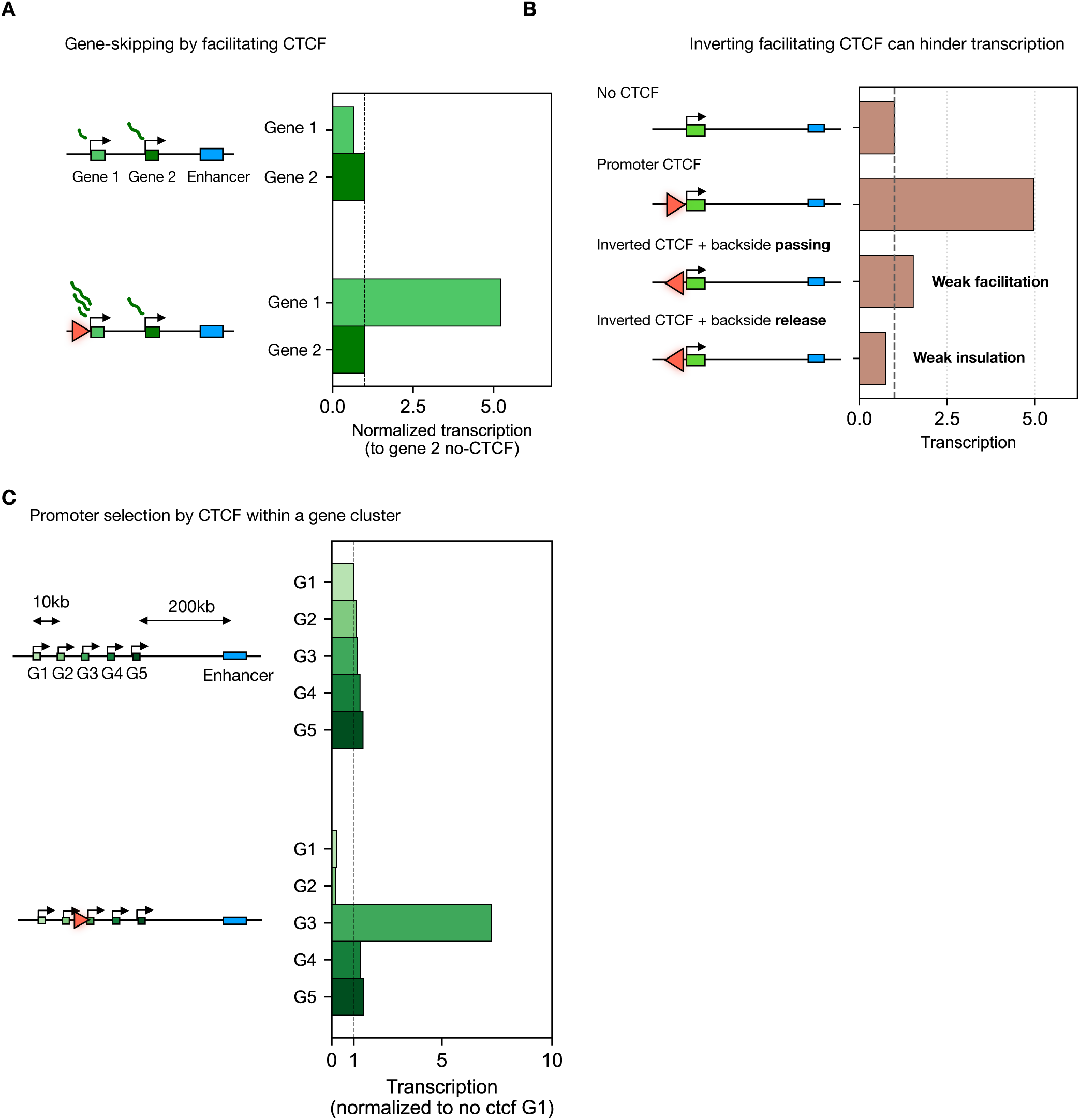
Predicted regulatory behaviors arising from CTCF modulation of cohesin enhancer-promoter bridging. **(A) *Promoter skipping by a facilitating CTCF site.*** In a two-promoter locus with a distal enhancer, the enhancer preferentially activates the proximal promoter in the absence of CTCF, as cohesin bridging is more frequent at shorter distances. Insertion of a CTCF site upstream of the distal promoter facilitates its activation by the enhancer, to levels that can exceed activation of the proximal promoter. Promoter competition (not modeled here) may further repress the proximal promoter. **(B) *Orientation-dependent facilitation***. Inversion of a facilitating promoter-proximal CTCF site is expected to reduce transcription. In the backside-passing regime, some facilitation persists due to occasional stalling of backside-approaching cohesins when the site is already occupied by a frontside-blocked cohesin. In the backside-release regime, expression drops below baseline because backside-encountering cohesins release the loop, actively disrupting enhancer-promoter bridges. **(C) *Promoter selection within a cluster.*** In a locus containing five closely spaced promoters (G1–G5) and a distal enhancer, insertion of a CTCF site just upstream of G3 selectively facilitates enhancer action on G3, while at the same time blocking cohesin bridges from reaching upstream promoters (G1-G2) and suppressing them. In this context, the inter-promoter spacing must be larger than *s\** to avoid spurious activation of neighboring promoters by facilitation.

**Figure S17.**
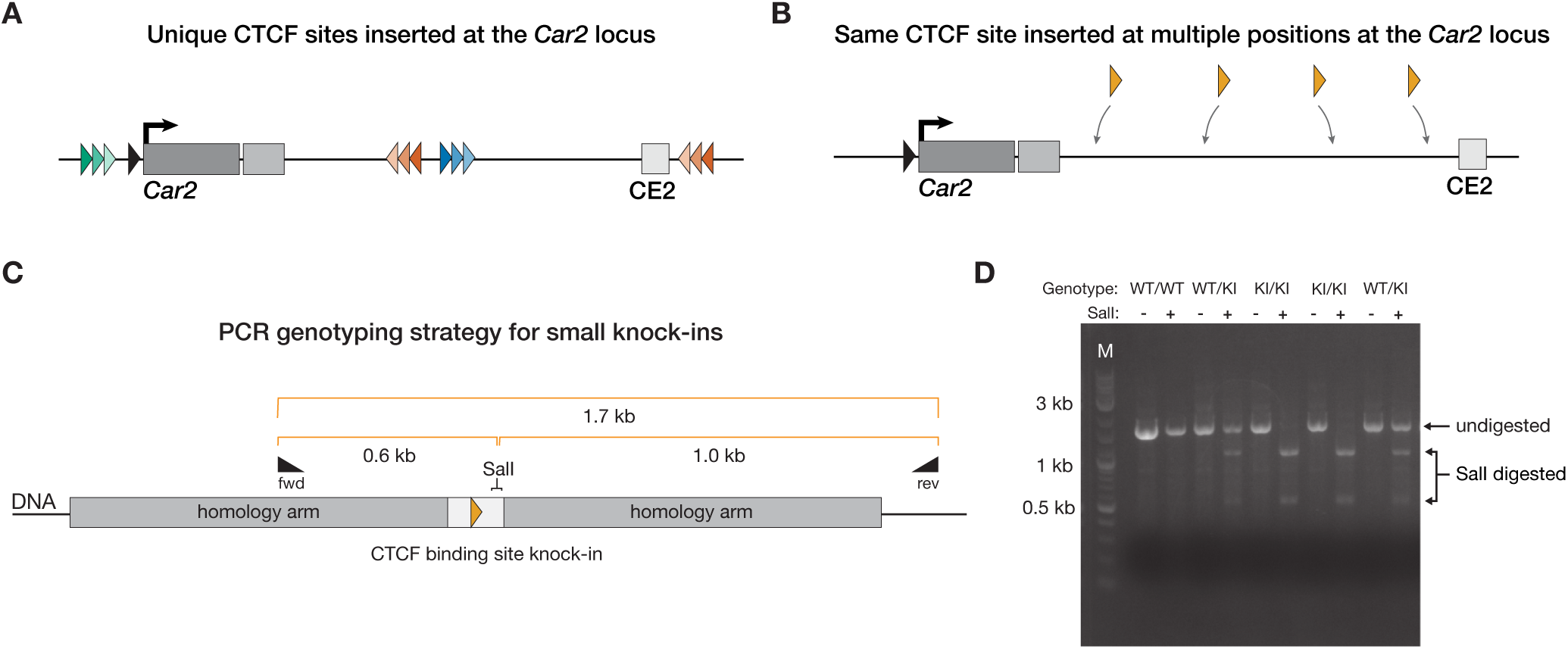
Details of CTCF sites used for insertions at the *Car2* locus and genotyping strategies. (A) Detailed schematic of the CTCF site cassette insertions at the *Car2* locus used in Figs. 3 to 5 (black: endogenous CTCF site). The sequence motif of each of the CTCF sites in the 3X cassettes was distinct from one another. The 3X cassettes in the reverse orientation (both intervening and at CE2) were identical to one another. See Methods for more details. (B) Detailed schematic of the single CTCF site inserted across the *Car2* locus, using the same motif sequence for each insertion (Fig. 6). See Methods for more details. **(C-D)** Strategy (C) and example gel (D) for genotyping small insertions at the *Car2* locus. Briefly, a unique restriction site was introduced with the CTCF site cassette that was used to digest a PCR product spanning the knock-in position. Desired heterozygous cell lines have both an upper, undigested WT band and lower, digested knock-in bands. DNA ladder marker (M) from NEB (cat# N3200). See Methods for more details.

**Video S1.**
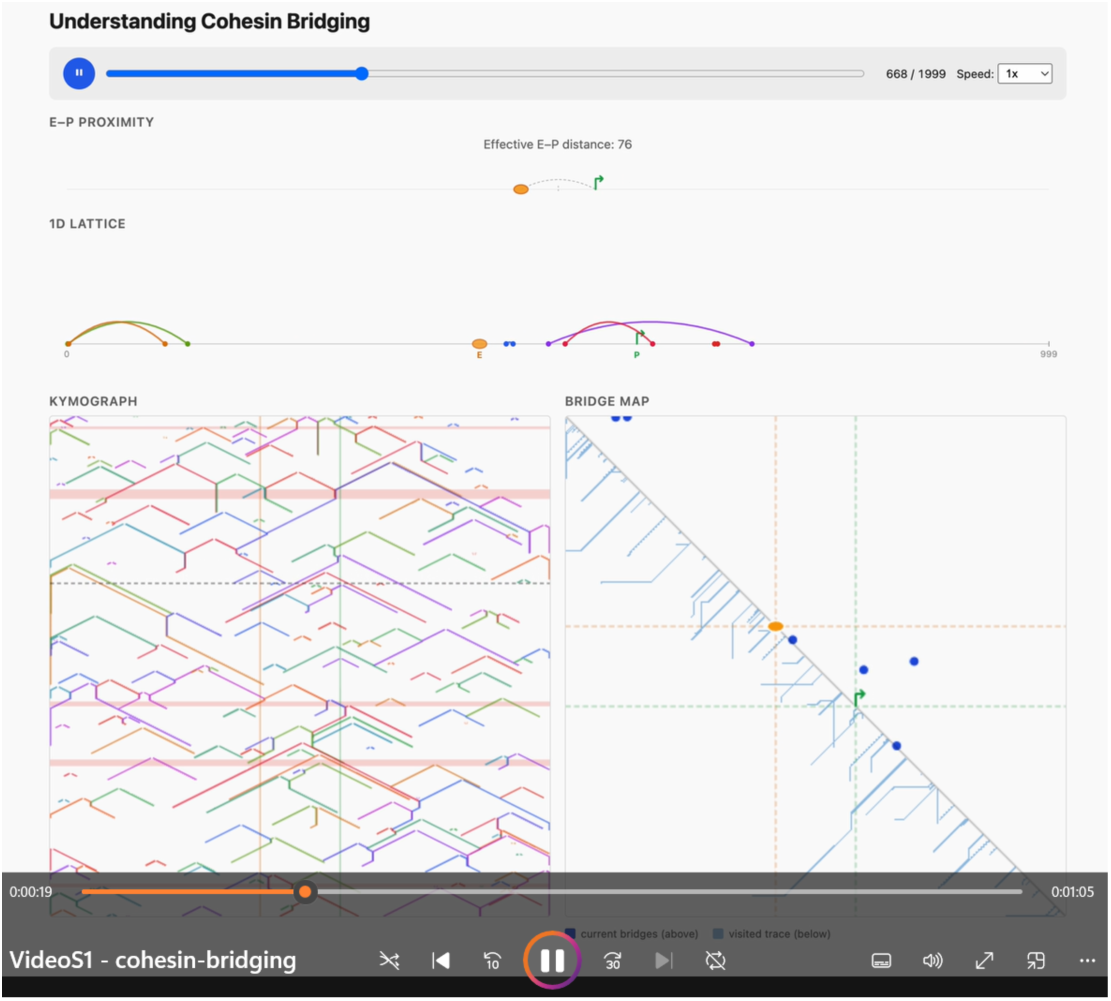
(cohesin-bridging.mp4). Visualization of enhancer-promoter activation by dynamic cohesin bridges. The video illustrates key features of the cohesin bridging model through multiple linked visualizations of simulated trajectories. Top: effective genomic distance (*s*eff) between the enhancer and promoter over time. Middle: positions of all cohesin loops along the locus. Bottom left: kymograph showing the history of cohesin loop trajectories. Bottom right: split contact map, with the upper-right triangle showing current extruder positions and the lower-left triangle showing the history of past trajectories. After an overview of cohesin dynamics, the video highlights individual enhancer-promoter bridging events, illustrating how single-loop and multi-loop configurations can contribute to transcriptional activation. Parameters: *λ* = 110 kb (cohesin processivity); *d* = 150 kb (average separation between cohesins); *L* = 160 kb (enhancer-promoter genomic distance); *s\** = 15 kb (activation distance threshold); symmetric two-sided extrusion for clarity of visualization. See online viewer at https://abdenlab.org/cohesin-bridging/

