## Supplementary text for "Cohesin bridging as a physical principle of enhancer-promoter communication"

### Supplementary text: Cohesin bridging as a physical principle of enhancer-promoter communication

### Part I

### Analytical theory for the dynamics of cohesin-bridged contacts in the absence of extrusion barriers.

Our goal is to compute the frequency of contacts between two loci separated by a distance L that arise from cohesin-bridging for two-sided extrusion and in the absence of barriers. In general, cohesin–cohesin interactions complicate this problem: collisions can cause stalling, reducing effective extrusion speed, while cooperative interactions can generate multi-loop contacts that extend bridging range. These effects are explored in the main text using simulations of interacting cohesins. Here, instead, we consider a simplified non-interacting model in which cohesins neither stall upon collision nor form multi-loop contacts. In this regime, cohesins act independently, and contact formation can be described by the probability that a single cohesin bridges the two loci. Although simplified, this framework is still informative because it isolates the core geometric and kinetic factors governing contact formation, thereby providing intuition for the full interacting system.

#### 1 Model framework and definitions

##### 1.1 Model description

We model DNA as a continuous one-dimensional line containing two target regions corresponding to the enhancer and promoter, of lengths $l_e$ and $l_p$, respectively, separated by a genomic distance $L$. Cohesins are represented by two points on this line corresponding to their anchoring sites of the molecule on DNA (basis of the loop). We consider cohesins bind uniformly along DNA at a rate per unit time per unit length $\kappa_b$. Once bound, a cohesin extrudes a loop at velocity $v$, until it dissociates (the loop grows at velocity $v$, each leg translocates at velocity $v/2$). Unbinding is assumed to occur at a constant rate $r_u$.

##### 1.2 Definition of a cohesin-bridged contact

Within this framework, we define a cohesin-bridged contact as an event in which cohesin brings the enhancer and promoter into close genomic proximity. Operationally, a contact occurs whenever the genomic separation between the enhancer and promoter is reduced below a threshold distance $s^*$ as a result of loop extrusion. In the full interacting system, contacts can arise from multi-loop configurations and are therefore defined more generally through the

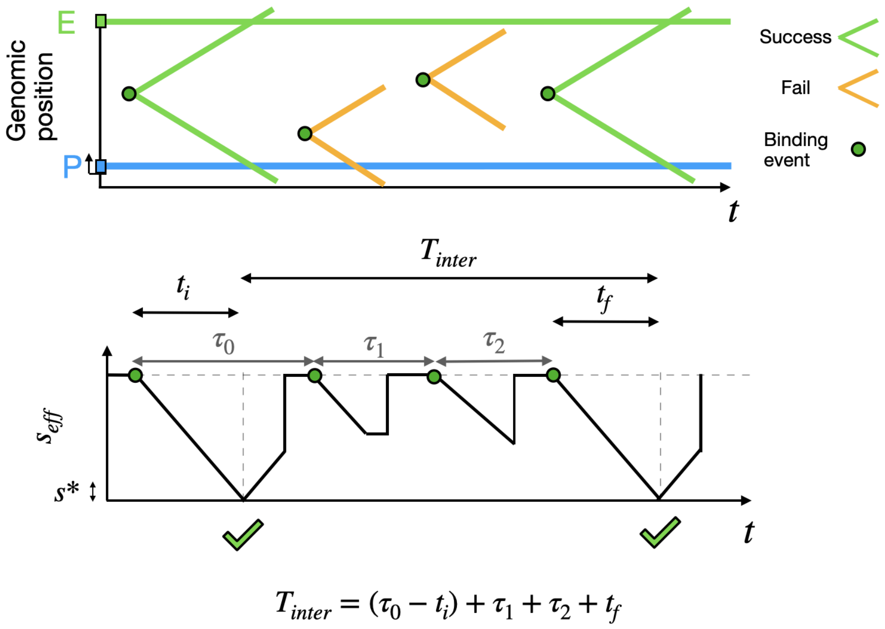

Figure 1: Top: Kymograph of cohesin trajectories. Cohesins that successfully bridge the enhancer and promoter are shown in green, and unsuccessful ones in orange. Bottom: Corresponding effective enhancer–promoter distance for the trajectories shown above.

effective 1D distance. Here, by contrast, we restrict our attention to single-loop events, so a contact corresponds specifically to a single cohesin, reducing the enhancer–promoter separation below $s^*$.

### 2 Contact statistics and first-passage formulation

#### 2.1 Contact statistics and inter-contact times

Let's compute the distribution of times between successive contacts, which we denote $T_{inter}$. To compute this quantity, we note that each cohesin loading event within the stretch of DNA separating the EP initiates a "trial" which can either result in a bridging event (success) or not (fail). Let $r_b$ be the total loading rate in the region separating the enhancer and promoter, so that the mean time between two "trials" is $\tau_b = \frac{1}{r_b}$.

In the case of non-interacting cohesins, we can derive a general formula to compute this quantity.

$$\langle T_{inter} \rangle = \frac{\tau_b}{p} \tag{1}$$

Where p denotes the probability that a single trial succeeds. In words, equation 5 says that the expected time to form a contact is simply the average waiting time between cohesin-binding events multiplied by the expected number of attempts required for one success. For example, if cohesin binds once per hour and only half of these binding events bring the loci close enough, then the mean time to form a contact is 2 hours.

Equation 5 is very general and can accommodate virtually any cohesin dynamics, barrier layouts, position-dependent velocities, binding and unbinding, and so on. The main challenge, however, lies in determining $p$. By the law of total probability, a general expression for $p$ can be written as an average over all possible loading sites. Since cohesins bind at random positions along the DNA, and their likelihood of successfully bridging depends on where they load, we have:

$$p = \int_0^L dx\, P(\text{success} \,|\, \text{loading at x}) P(\text{loading at x}) \tag{2}$$

From this simple formula, we can compute the frequency of contacts for various situations—including configurations with CTCF barriers, different EP distances, and varying enhancer/promoter lengths. For completeness, we provide a short derivation below.

**Derivation of Eq. 5**

##### 2.1.1 Demonstration of equation 5

To demonstrate equation 5, we need to write $T_{inter}$ as a function of the time between cohesin binding events. The following formula can be trivially derived from the diagram in Figure 1:

$$T_{inter} = (\tau_0 - t_0) + \tau_1 + ... + \tau_k + t_k \tag{3}$$

Where $\tau_i$ denotes the time between binding i and i+1 and $t_i$ and $t_f$ are the times for extruder 0 (resp. k) to reach the EP after binding.
Because the lifetimes of different extruders are independent and identically distributed, $t_0$ and $t_k$ have the same mean, such that $\langle t_0 - t_k \rangle = 0$ Therefore, the expected inter-contact time reduces to

$$\langle T \rangle = \langle \tau_0 + \ldots + \tau_{k-1} \rangle \tag{4}$$

Each $\tau_i$ is exponentially distributed with mean $\langle \tau \rangle = 1/r_b$, and the number of trials $k$ is geometrically distributed with mean $\langle k \rangle = 1/p$, where $p$ is the success probability of a single trial. Using the law of total expectation, we obtain

$$\langle T_{inter} \rangle = \langle k \rangle \langle \tau_b \rangle = \frac{\tau_b}{p} \tag{5}$$

### 3 Contact frequency for an isolated enhancer and promoter

We now compute the mean inter-contact time $\langle T_{inter} \rangle$ for an enhancer–promoter pair of finite lengths, allowing for a finite slack $s^*$. Because the full calculation is somewhat involved, we decompose it into two simpler subproblems. We first consider finite-size elements without slack ($s^* = 0$), and then treat point-like elements with an allowed slack ($l_e = l_p = 0$). The general result follows by combining these two limits.

#### 3.1 Contact frequency for a point-like E–P pair

##### 3.1.1 Cohesin binding outside the antenna cannot bridge the E–P pair

We consider a point-like enhancer and promoter under symmetric extrusion. If cohesin loads at a distance $x$ from the midpoint between E and P, the smallest separation it can achieve between the two loci is $2|x|$ (2).

A successful contact therefore, requires

$$2|x| \leq s^*.$$

This defines a central region of size $s^*$ around the midpoint from which loading can produce a contact. We refer to this region as the *antenna.* Cohesin loading outside the antenna cannot generate a contact.

This can also be visualized on a contact map. In this representation, each cohesin is described by the genomic positions of its two anchoring points (the base of the loop), which correspond to a point in the contact map. A contact occurs when these two points fall within a region where the enhancer and promoter are brought into proximity. We therefore define a contact zone as the set of positions in the contact map for which the presence of a cohesin results in an enhancer–promoter contact. In the limit of a point-like enhancer and promoter with finite slack, this contact zone takes the shape of a square. (Fig. fig. 3).

In the case of symmetric extrusion, each cohesin follows a straight-line trajectory in the contact map. The antenna can therefore be defined as the set of loading positions whose trajectories intersect the contact zone (See fig. 3)).

##### 3.1.2 Cohesin lifetime on DNA is exponentially distributed

Cohesins that bind inside the antenna must extrude far enough to reach both the enhancer and the promoter. The probability of doing so depends on the

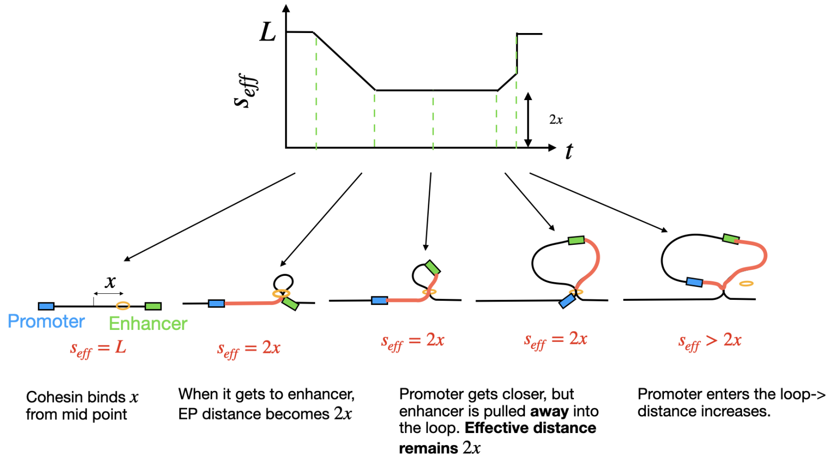

Figure 2: Schematic of the evolution of the effective enhancer–promoter distance during a loop extrusion event.

unbinding rate, extrusion velocity, and the enhancer–promoter separation $L$.

Assuming a constant unbinding rate $r_u$, the lifetime of a bound cohesin follows an exponential survival law. The probability that an extruder remains bound longer than time $T$ is

$$P(t > T) = e^{-r_u T}.$$

Because extrusion is assumed to proceed at a constant deterministic velocity $v$, the extruded length and lifetime are related by

$$l = vt.$$

Thus, the probability of extruding at least a length $l$ is

$$P(l > L) = e^{-r_u L/v} = e^{-L/\lambda},$$

where $\lambda = v/r_u$ is the processivity, i.e., the mean extruded length.

For a cohesin loaded within the antenna to produce a contact, it must extrude a genomic length $L - s^*$.

###### 3.1.3 Computing $\langle T_{inter} \rangle$

We now compute the success probability $p$ using Eq. (2):

$$p = \int_0^L dx \, P(\text{success} \mid x) \, P(\text{loading at } x). \tag{6}$$

From the considerations above, a cohesin loaded inside the antenna must extrude a length $L - s^*$ to generate a contact, giving a success probability $e^{-(L-s^*)/\lambda}$. Outside the antenna, the success probability is zero:

$$P(\text{success} \mid x) = \begin{cases} e^{-(L-s^*)/\lambda} & \text{if } x \in (-s^*/2, s^*/2) \\ 0 & \text{otherwise.} \end{cases} \tag{7}$$

In this point-like limit, the required extrusion length does not depend on the exact loading position within the antenna. (For finite-size elements, the required distance will depend on $x$.)

Assuming homogeneous loading,

$$P(\text{loading at } x) = 1/L.$$

Therefore,

$$p = \frac{1}{L} \int_{-s^*/2}^{s^*/2} dx\, e^{-(L-s^*)/\lambda} \tag{8}$$

$$= \frac{s^*}{L} e^{-(L-s^*)/\lambda}. \tag{9}$$

Finally, using $r_b = \kappa_b L$ and $\langle T_{inter} \rangle = 1/(r_b p)$,

$$\langle T_{inter} \rangle = \frac{1}{\kappa_b s^*}\, e^{(L-s^*)/\lambda}. \tag{10}$$

This result has a simple interpretation: $\kappa_b s^*$ is the rate of cohesin loading within the antenna, while the exponential factor reflects the probability that cohesin remains bound long enough to extrude the required distance. Consequently, the contact rate decreases exponentially with characteristic scale $\lambda$.

##### 3.2 finite length enhancer-promoter pair

We now repeat the calculation of $\langle T_{inter} \rangle$ for an enhancer–promoter pair with finite lengths $l_e$ and $l_p$, and no slack. A contact is defined as a configuration in which cohesin simultaneously contacts both elements.

As before, we compute the success probability $p$ from Eq. (2). Under symmetric extrusion, successful bridging requires cohesin to load in the central region between the two elements. For finite-length elements, the antenna has size $(l_e + l_p)/2$ (Fig. 4).

Unlike the point-like case, the extrusion distance now depends on the loading position $x$: cohesins loaded closer to the midpoint have a shorter distance to extrude and are therefore more likely to succeed. The success probability conditional on loading at position $x$ is:

$$P(\text{success} \mid x) = \begin{cases} e^{-(L+2|x|)/\lambda} & x \in [-l_e/2,\, l_p/2] \\ 0 & \text{otherwise,} \end{cases} \tag{11}$$

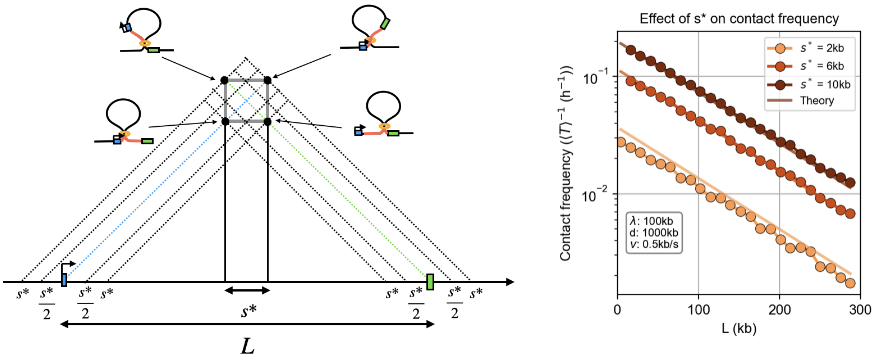

Figure 3: (Left) The contact zone (grey square) defines all cohesin configurations that bring the enhancer and promoter within a distance s*. Under symmetric extrusion, cohesins follow straight-line trajectories, and only those loading within the antenna (set of trajectories intersecting the contact zone) can generate contacts. (Right) Contact frequency as a function of enhancer–promoter distance L for different values of s*. Symbols: simulations; lines: analytical prediction.

where $\lambda = v/r_u$ is the loop-size processivity.
Assuming homogeneous loading in the E–P interval,

$$p = \frac{1}{L}\int_{-l_e/2}^{l_p/2} e^{-(L+2|x|)/\lambda}\, dx \tag{12}$$

$$= \frac{\lambda}{2L} e^{-L/\lambda}\Big[2 - \big(e^{-l_e/\lambda} + e^{-l_p/\lambda}\big)\Big]. \tag{13}$$

Using $\langle T_{inter}\rangle^{-1} = r_b p$, we obtain

$$\langle T_{inter}\rangle^{-1} = r_b \left(\frac{\lambda}{2}\right) e^{-L/\lambda}\Big[2 - \big(e^{-l_p/\lambda} + e^{-l_e/\lambda}\big)\Big]. \tag{14}$$

In the regime $\lambda \gg l_e, l_p$, expanding to first order yields

$$\boxed{\langle T_{inter}\rangle^{-1} \simeq r_b \frac{l_p + l_e}{2} e^{-L/\lambda}.} \tag{15}$$

This form has a simple interpretation: $r_b(l_p + l_e)/2$ is the effective loading rate within the antenna, and $e^{-L/\lambda}$ is the probability of extruding a loop spanning the E–P distance. The full expression accounts for the additional distance that off-center cohesins must extrude.

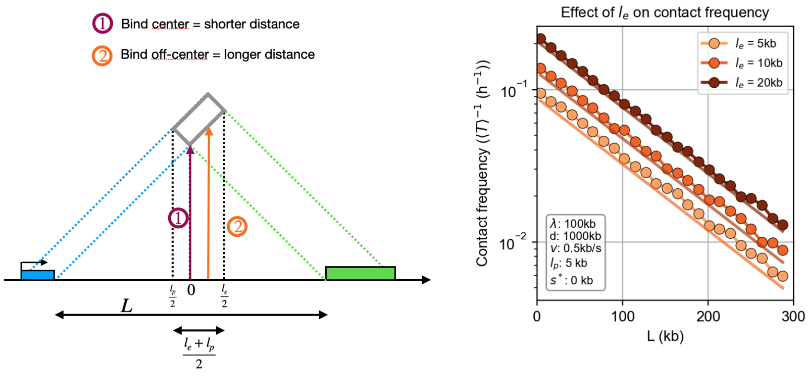

Figure 4: Effect of enhancer or promtoer length on contact frequency. (Left) Schematic illustrating how the loading position of cohesin within the enhancer–promoter interval affects the required extrusion distance to the contact zone (grey). Cohesins binding near the center (1) require shorter extrusion distances to bridge the enhancer and promoter, while off-center loading (2) requires longer extrusion, reducing the probability of successful contact slightly. (Right) Contact frequency as a function of enhancer–promoter distance L for different enhancer lengths $l_e$. Increasing $l_e$ expands the antenna and increases contact frequency. Symbols: simulations; lines: analytical prediction.

##### 3.3 Finite-length enhancer-promoter pair with slack

We now consider the full problem in which the enhancer and promoter have finite lengths $l_e$ and $l_p$, and activation occurs when the effective E–P distance falls below $s^*$.

The antenna is enlarged relative to the previous limits: cohesin can load within a central slack region of width $s^*$, and can additionally load further toward either element by up to $l_e/2$ (enhancer side) or $l_p/2$ (promoter side), at the cost of requiring a larger loop. Formally, the position-dependent writes:

$$P(\text{success} \mid x) = \begin{cases} \exp\left(-\dfrac{L-s^*}{\lambda}\right), & |x| \leq \dfrac{s^*}{2}, \\ \exp\left(-\dfrac{L+2|x|-2s^*}{\lambda}\right), & \dfrac{s^*}{2} < |x| \leq \dfrac{l_e+s^*}{2} \text{ and } x \leq 0 \text{ or } \dfrac{s^*}{2} < |x| \leq \dfrac{l_p+s^*}{2} \text{ and } x \geq 0, \\ 0, & \text{otherwise.} \end{cases} \tag{16}$$

Averaging the position-dependent success probability over the antenna to compute $p$, and injecting $p$ into (5) yields:

$$\langle T_{inter}\rangle^{-1} = r_b \, \frac{\lambda}{2L} \, e^{-(L-s^*)/\lambda} \left[2 - e^{-(l_e+s^*)/\lambda} - e^{-(l_p+s^*)/\lambda}\right]. \tag{17}$$

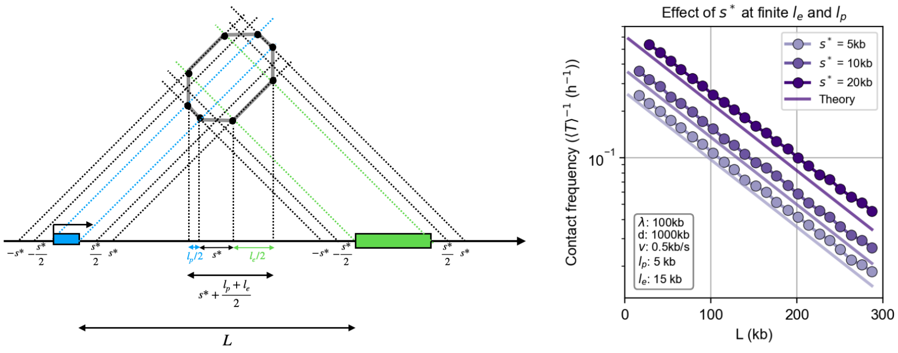

Figure 5: Same as 3 and 4 but for a finite-sized enhancer and promoter plus allowed slack.

$$\langle T_{inter} \rangle^{-1} = r_b \, \frac{\lambda}{2L} \, e^{-(L-s^*)/\lambda} \left[ 2 - e^{-(l_e+s^*)/\lambda} - e^{-(l_p+s^*)/\lambda} \right] \qquad (18)$$

*This expression is used to generate the theoretical predictions shown in the main Fig. 2A.* In the experimentally relevant regime $\lambda \gg l_e, l_p, s^*$, it simplifies to

$$\boxed{\langle T_{inter} \rangle^{-1} \simeq r_b \left( s^* + \frac{l_e + l_p}{2} \right) e^{-L/\lambda}.} \qquad (19)$$

Thus, the contact frequency is simply the rate of loading within the antenna multiplied by the probability of surviving long enough to span the E-P distance.
